# Topologically associated domains are ancient features that coincide with Metazoan clusters of extreme noncoding conservation

**DOI:** 10.1101/042952

**Authors:** Nathan Harmston, Elizabeth Ing-Simmons, Ge Tan, Malcolm Perry, Matthias Merkenschlager, Boris Lenhard

## Abstract

In vertebrates and other Metazoa, developmental genes are found surrounded by dense clusters of highly conserved noncoding elements (CNEs). CNEs exhibit extreme levels of sequence conservation of unexplained origin, with many acting as long-range enhancers during development. Clusters of CNEs, termed genomic regulatory blocks (GRBs), define the span of regulatory interactions for many important developmental regulators. The function and genomic distribution of these elements close to important regulatory genes raises the question of how they relate to the 3D conformation of these loci. We show that GRBs, defined using clusters of CNEs, coincide strongly with the patterns of topological organisation in metazoan genomes, predicting the boundaries of topologically associating domains (TADs) at hundreds of loci. The set of TADs that are associated with high levels of non-coding conservation exhibit distinct properties compared to TADs called in chromosomal regions devoid of extreme non-coding conservation. The correspondence between GRBs and TADs suggests that TADs around developmental genes are ancient, slowly evolving genomic structures, many of which have had conserved spans for hundreds of millions of years. This relationship also explains the difference in TAD numbers and sizes between genomes. While the close correspondence between extreme conservation and the boundaries of this subset of TADs does not reveal the mechanism leading to the conservation of these elements, it provides a functional framework for studying the role of TADs in long-range transcriptional regulation.

## Introduction

In Metazoa, many genes involved in developmental regulation are surrounded by syntenic arrays of conserved non-coding elements (CNEs) [1, 2, 3, 4, 5]. These CNEs exhibit extreme levels of conservation over a large number of base pairs, in some cases more than the equivalent conservation of protein-coding regions [6]. Several studies have shown that individual CNEs can act as transcriptional enhancers to drive complex spatiotemporal expression patterns [7, 8, 9]. However, no known source of selective pressure is able to account for their extreme levels of conservation [10].

The expression patterns driven by CNEs overlap with those of nearby developmental regulators, suggesting that they act as regulatory elements for these genes [11, 12, 13, 14, 15, 9]. However, overlap is often partial, with different CNEs driving expression in different spatiotemporal subdomains [16, 7, 8], suggesting that multiple regulatory elements are required to drive complex expression patterns. The syntenic organisation of clusters of CNEs, called genomic regulatory blocks (GRBs) [17, 3], supports the idea that they are ensembles of regulatory elements that regulate important developmental genes. The syntenic organisation between CNEs and their target genes results from the necessity of keeping regulatory elements, which may be separated from their target genes by large genomic distances, in cis with the gene under long-range regulation [18, 19]. In addition to this target gene (or genes in the case of gene clusters) under developmental regulation, a GRB can harbour several other genes that are not detectably regulated by these elements (bystander genes). Target and bystander genes differ with respect to their promoter structure, patterns of epigenetic modification and range of biological functions [3, 20, 21].

Long-range regulation depends on the interaction of a target gene promoter with enhancers located at up to megabase distances, which need to be brought into close physical proximity in the nucleus. Insights into interactions at this scale and their roles in development and differentiation have been provided by the development of chromatin conformation capture methods [22]. Interactions between regulatory elements located within the introns of bystander genes and the promoters of target genes have been identified using 3C [23]. The 3D structure of vertebrate Iroquois clusters, which are known GRBs, appears to be highly conserved across vertebrates [24], and is thought to result from enhancer sharing and co-regulation of members of the cluster during development. Chromatin interactions can also be investigated genome-wide using Hi-C [25]. This has revealed regions of the genome which preferentially self-interact, known as topologically associating domains (TADs) or contact domains [26, 27]. Regulatory elements and genes preferentially interact within the same TAD, suggesting that the boundaries of TADs may act to restrict the influence of enhancers [28, 29, 30].

Despite containing cell-type specific promoter-enhancer interactions, the structure of TADs in mammals is remarkably invariant across different cell types [26, 27, 31], including sperm [32], and between species [26, 33]. TADs have previously been found to correspond with other large-scale genomic features, including replication domains (RD) [34, 35, 36, 37], lamina-associated domains (LADs) [36] and Polycomb-repressed domains [38], and to have boundaries that coincide with conserved CTCF binding [33]. While it appears that human, mouse and *Drosophila* chromosomes are segmented into TADs along their entire length [26, 38], in *C. elegans* their occurrence varies both between and along different chromosomes [39]. Plant chromosomes do not seem to be organised into TADs, except for isolated TAD-like structures at a limited set of loci [40]. At longer length scales, Hi-C data suggests that TADs are organised into two major compartments, which correspond to open (compartment A) or closed chromatin (compartment B), which tend to self-associate within the nucleus [25]. TADs may switch compartment depending on the activity of genes within them [31], therefore it has been suggested that these form the regulatory units of the genome [41].

The stability of TADs, the variation in their size and their ubiquitous presence across different metazoan phyla lead us to hypothesise that they might represent evolutionarily stable features of loci involved in developmental regulation. Given that our previous work strongly suggested that genomic regulatory blocks (GRBs) correspond to the regulatory domains of key developmental genes [3, 17, 7], and that little is known about their relationship with genome-wide 3D organisation, we set out to explore the relationship between GRBs and TADs systematically in both vertebrates and invertebrates.

## Results

In this paper, we define a CNE as a non-coding element that has a high percentage identity over a defined number of base pairs in a comparison between two species (see Methods). We operationally define GRBs as discrete regions of the genome with a high density of syntenic CNEs. The locations of putative GRBs and their approximate spans can be visualised by plotting the density of CNEs in a sliding window (Fig. 1A). We developed a CNE clustering approach that robustly estimates the extent of GRBs, based solely on the distribution of syntenic CNEs in the genome (see Methods). The procedure is shown schematically in Supplementary Fig. 1. This allowed the examination of the physical extent and boundaries of putative GRBs, and their comparison with higher-order chromatin organisation.

**Figure 1:**
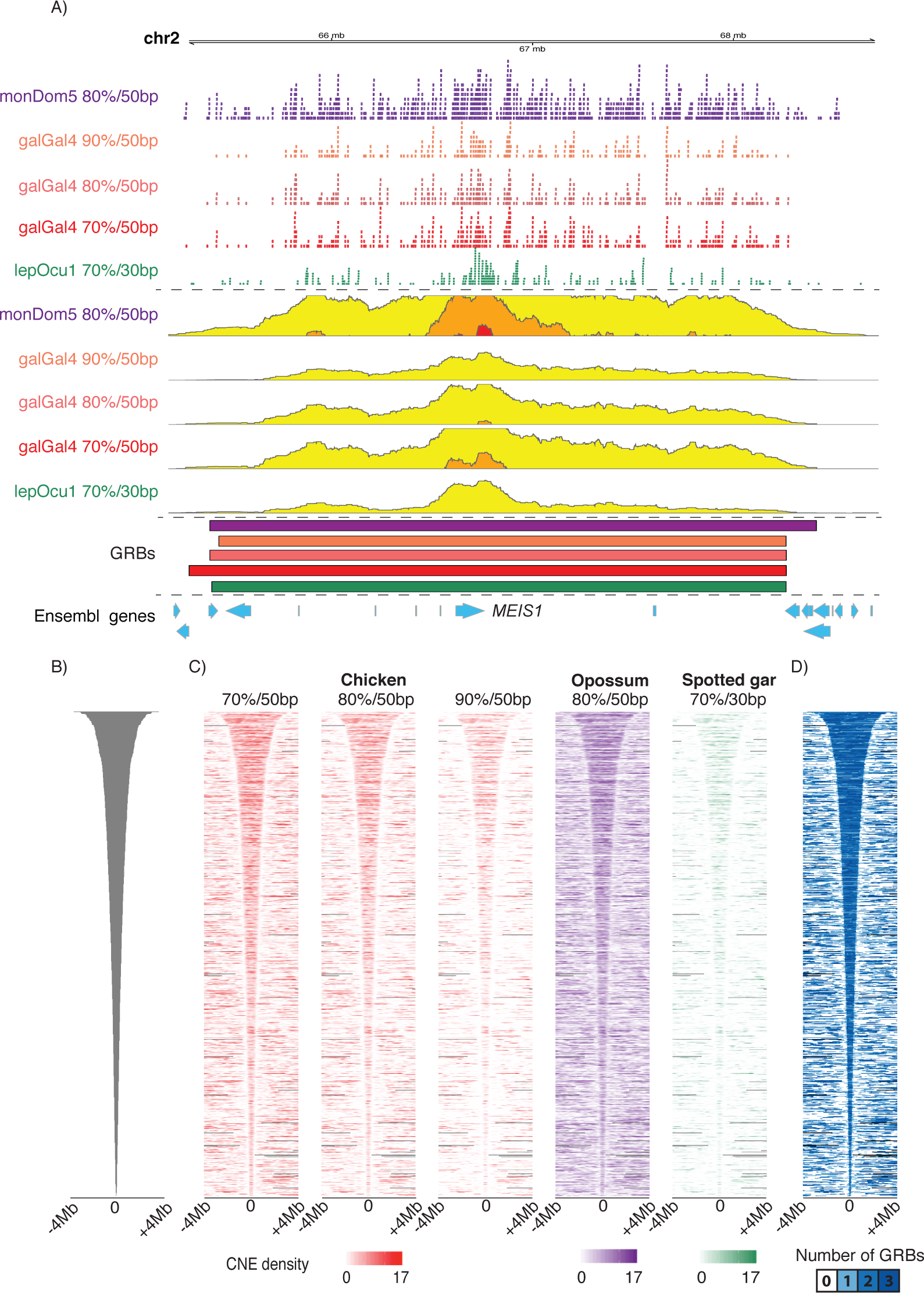
The boundaries of GRBs are highly consistent regardless of the thresholds or species involved. A) The human *MEIS1* locus is spanned by arrays of conserved noncoding elements identified in comparisons with opossum, chicken and spotted gar. These CNEs can be visualised as a smoothed density, shown here as a horizon plot. The boundaries of the proposed GRBs at the *MEIS1* locus are highly consistent regardless of the species or thresholds involved. B) All hg19-galGal4 GRBs centred and ordered by length of the GRB C) Distribution of CNEs in a window of 8Mb around the centre of the hg19-galGal4 GRBs for different sets of CNEs. D) Overlap of GRBs obtained from predictions using hg19-monDom5, hg19-galGal4, hg19-lepOcu1.

Hi-C datasets for various species and cell types were obtained [26, 38, 31] and processed to generate genome-wide interaction maps (see Methods). TADs are usually visualised as triangular regions of increased signal in interaction matrices. Their span is typically identified by calculating a directionality index for each region in the genome that quantifies its level of upstream or downstream interaction bias, which is then processed to segment the genome into a discrete set of TADs for downstream analysis [42, 26, 43, 39].

## GRBs show conserved boundaries robust to chosen thresholds and evolutionary distance

The development of a CNE clustering method allowed the generation of sets of GRBs between species at various evolutionary distances. GRBs were identified in the human genome using the distribution of CNEs observed in comparisons with opossum (160 Mya separation), chicken (320 Mya), and spotted gar (430 Mya) [44]. GRBs between human and chicken (hg19-galGal4) were identified using a number of thresholds: 819 GRBs were identified using CNEs showing 70% identity over 50bps (Fig. 1B), 672 using 80% over 50bps and 468 using 90% over 50bps. Regardless of the threshold used to identify CNEs present between human and chicken, the density of CNEs and associated GRB predictions remain highly concordant (Fig. 1 and Fig. S2). 1160 GRBs were predicted between human and opossum (hg19-monDom5) using CNEs showing 80% identity over 50bps and 719 between human and spotted gar (hg19-lepOcu1) using CNEs showing 70% identity over 30bps.

As evolutionarily conserved features, we expect the generated GRBs to have stable boundaries. Indeed, the span of GRBs identified using hg19-galGal4 show close agreement with the distribution of CNEs identified from hg19-monDom5 and hg19-lepOcu1 comparisons (Fig. 1C). Examination of a number of GRBs revealed a marked correspondence of their boundaries between evolutionarily distant species. Comparison of the boundaries of hg19-galGal4 GRBs that overlapped with individual hg19-monDom5 GRBs found that 276 (50% - 554 in total) had boundaries that differed by less than 150kb. A subset of hg19-galGal4 GRBs appear to have similar boundaries to hg19-monDom5 and hg19-lepOcu1 GRBs (Fig. 1D), including GRBs containing well-known developmental regulators, including *MEIS1* and *IRX3* (Fig. 1A, Fig. S2). This suggests that the span of several GRBs is invariant to the evolutionarily distances involved in identifying them.

Two main problems can occur with the prediction of GRB span from CNE density. If the synteny between two adjacent GRBs is conserved in the species used for comparison and the GRBs are extremely close together, there is no information from the CNE density alone that would enable their separation during clustering. At several loci, separate GRBs are known to border each other, e.g. *TOX3* / *SALL1* [13, 45] (Fig. S2A) and *PAX6* / *WT1* [7] (Fig. S2D). For that reason, we expect to have a fraction of estimated GRB spans that contain multiple adjacent GRBs, and predict that this problem will be more prevalent with the longest span predictions (i.e. the top set of predictions in Fig. 1B). Alternatively, since the density of CNEs along a GRB is non-homogeneous, a single GRB can be split into two or more putative regions. This may occur if there is a sparsely populated region of CNEs between two dense regions, or if the distance between two putative regions is large. We expect this problem will be more prevalent in predictions of smaller GRBs (i.e. the lower set of predictions in Fig. 1B) and be responsible for the majority of poor predictions generated by our method. The phylogenetic distance and conservation thresholds are able to manage some of these outcomes.

The identification of GRBs and the concordance of their boundaries over multiple species and thresholds suggests that our clustering method is robust and that GRBs represent evolutionarily conserved structures. For our human-centric comparisons we used GRBs identified using CNEs conserved between human and chicken at 70% identity over 50bp, which provide the best compromise between GRB coverage and the ability to separate adjacent GRBs.

## GRB boundaries strongly correlate with the boundaries of TADs

To visualise the relationship between the identified GRBs and TADs, we produced heatmaps of genomic regions centred on the GRB and ordered by GRB size, in which the GRBs and any features that correlate with them show a characteristic funnel shape. To show the TAD data for the GRB regions shown on the heatmap, we used Hi-C directionality index (positive/red when this region is preferentially interacting with regions downstream, and negative/blue when this region is preferentially interacting with regions upstream; one TAD is typically a red region followed by similar-sized blue region). Examination of Hi-C directionality index of predicted GRB spans revealed a striking pattern of sharp changes in directionality index, typically found at TAD boundaries, at the boundaries of GRBs in both human (Fig. 2A, S3G) and *Drosophila melanogaster* (Fig. 2B). The largest of the predicted GRB spans show multiple changes of directionality within the region, in line with our expectation that these regions may represent multiple closely spaced GRBs. Also, the distinctive funnel shape is lost at the bottom of the plot, in line with predictions regarding the performance of our clustering approach. The association between GRBs and directionality index is present regardless of the cell line used (Fig. S3G).

**Figure 2:**
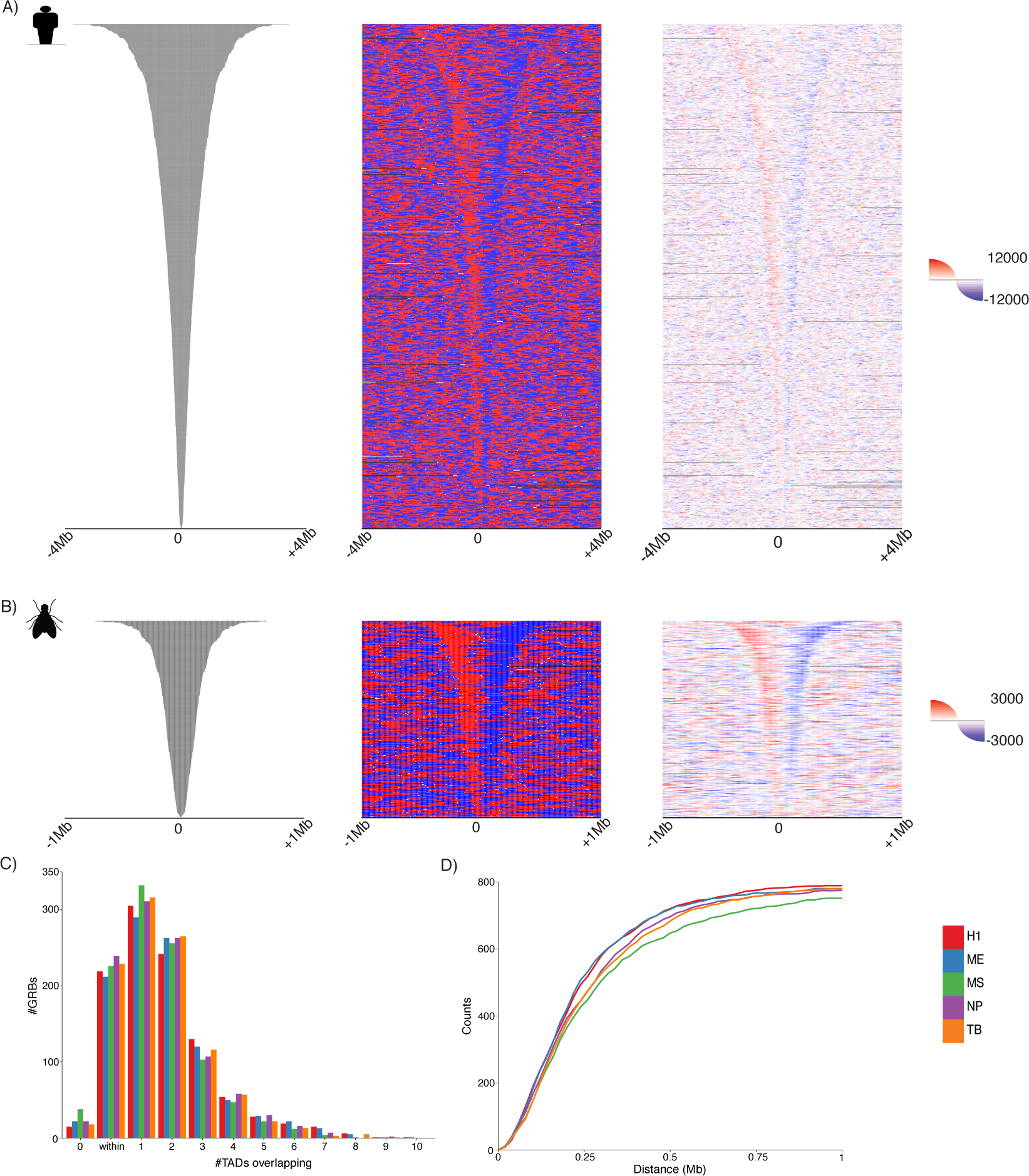
The boundaries of GRBs predict the boundaries of TADs in multiple evolutionarily distant species. A) Heatmaps representing H1-ESC directionality index spanning an 8 Mb window around the centre of putative hg19-galGal4 GRBs. Showing both the overall direction (middle panel, red for downstream, blue for upstream) and the average raw directionality score in 5kb bins (right panel). B) Heatmaps of *Drosophila* embryo Hi-C directionality index spanning a 2Mb window around the centre of dm3-droMoj3 GRBs. Showing both the overall direction (middle panel, red for downstream, blue for upstream) and the average raw directionality score in 1 kb bins (right panel). C) A large number of GRBs was found to be located within individual TADs (identified using HOMER) or overlapping only a single TAD, regardless of cell lineage (H1-ESC (H1), mesenchymal stem cells (MS), mesendoderm (ME), neural progenitor cells (NP) and trophoblast-like (TB)). D) Cumulative distribution of distance to nearest TAD (HOMER) boundaries from GRB boundaries in different cell lineages considering both edges i.e. both the start and end position of a GRB lie within X kb of the nearest TAD start and end.

To further explore the relationship between GRBs and topological domains, we compared the span of our sets of putative GRBs with TADs identified in several human cell lineages [31] and in *Drosophila* embryos [38]. Regardless of either the cell lineage or the method used to identify TADs, a large number of GRBs was found to be located within individual TADs or overlapping only a single TAD (Fig. 2C, Fig. S3A). An investigation of the distances between putative GRB boundaries and the nearest TAD boundary revealed a close association between them, with both edges of 236 GRBs (29%) lying within 120kb (i.e. three Hi-C bins, see Methods) of the nearest TAD boundary in H1-ESCs (Fig. 2D, Fig. S3B). For 369 GRBs (45%), at least one edge lies within 120kb of the nearest H1-embryonic cell TAD boundary (p < 1e-4, permutation test, Fig. S3E). This relationship between CNE density and topological structure was also observed in high-resolution Hi-C [27] generated in GM12878 cells (Fig. S4), with 398 GRBs (49%) having at least one edge within 50kb of the nearest outermost contact domain boundary (p < 1e-4, permutation test, Fig. S4C). We confirmed this association in an evolutionarily distant species using a set of GRBs identified between *Drosophila melanogaster* and *Drosophila mojavensis* (dm3-droMoj3) (63 Mya). 317 GRBs were identified using CNEs showing 96% identity over 50bp. Again, the majority of GRBs were located within or were overlapping with one TAD (Fig. S3C), with both edges of 110 GRBs lying within 60kb of a TAD boundary (p < 1e-4, permutation test, Fig. S3D). Therefore, the boundaries of GRBs identified using our method can predict at least one edge of a subset of TADs and accurately predict the location of both edges at one third of loci in both human and *Drosophila*.

Importantly, we show that the correspondence between the TADs and GRBs is not merely a consequence of the presence of clusters of enhancers in these regions: the spans of neither GRBs nor TADs coincide with the density of H3K27ac identified using ChIP-seq (Fig. S3H). The observed general depletion of H3K27ac signal within GRBs and an increase in signal at a small subset of boundaries are expected given that GRBs are typically associated with regions of low gene density and that H3K27ac is found at the promoters of transcriptionally active genes. This observation, along with the lack of predictive power in using H3K27ac (and other histone modifications) to predict the extent of TADs [46], imply that the potential enhancer activity of CNEs is not sufficient to explain the observed pattern. Therefore, there is a strong genome-wide correspondence between the genomic regions enriched for extreme noncoding conservation and those identified as TADs in Hi-C experiments. This relationship is robust with respect to cell lineage, species and the computational method used to identify TADs.

As predicted by our genome-wide analysis, individual loci of known target genes showed a strong concordance between CNE density and topological structure (Fig. 3 and S5). *MEIS1* is a transcription factor involved in multiple developmental processes [47] and disorders [15]. The *MEIS1* GRB and its topological domain (Fig. 3A) encompass all of the CNEs identified as enhancers in reporter assays [48], with a strong resemblance between GRB boundaries, which are highly concordant regardless of the species used to identify them (Fig. 1A), and the topological organisation of this locus as defined by Hi-C. A similar pattern is observed at the locus containing *RUNX2* (Fig. 3B), a transcription factor required for proper osteoblastic differentiation and bone development [49], with CNE density correlating with TADs at this locus. The boundaries of the *IRX3/5/6* GRB (Fig. 3C) are highly consistent regardless of the species used to identify CNEs (Fig. S2A), with this GRB being highly predictive of TAD boundaries, showing a strong concordance with both the directionality index and interaction matrix. Importantly, the TAD contains the *FTO* gene, whose intronic regulatory elements contact IRX genes as predicted by the GRB model [14] and confirmed experimentally [50, 51]. Our method identified the region containing *TOX3* and *SALL1* as a single GRB (Fig. 3C), however the boundary between the regulatory domains of these genes identified in enhancer screens [13] is reflected by a TAD boundary located in the middle of this region. The regulatory landscape of the *HoxD* cluster has previously been found to be better predicted by synteny than by its topological structure [52], indeed at this locus the span of the potential interactions closely resembles the distribution of CNEs at this locus (Fig. S5A).

It is known that CNEs cluster around orthologous transcription factors in both vertebrates and arthropods, however the elements themselves exhibit little sequence similarity between phyla [53, 3]. *Drosophila* loci known to contain important developmental genes show patterns of association between CNE density and topological organisation similar to those seen in human (e.g. *hth*, *pros*, *CG34114* (Fig. S5B) and the *Antennapedia* complex (Fig. S5C)). Vertebrate homologs of these genes (e.g. *MEIS1* / *MEIS2* (*hth*) and *PROX1* (*pros*),*HOX* (*Antp*),*SOX1/2/3* (*soxN*) are located in regions of extreme non-coding conservation (Fig. 1A, S1) in hg19-galGal4 comparisons, which are highly predictive of the span of interactions observed in Hi-C (Fig. 3). In addition, at the *Six* gene loci in the sea urchin *Strongylocentrotus purpuratus*, the regulatory landscapes defined by the span of promoter contacts identified using 4C [54] closely correlates with lineage-specific CNE density at this locus (Fig. S6A).

These results show overwhelming evidence for a strong concordance between CNE density and topological organisation in the genomes of vertebrates and arthropods; with evidence at one locus in echinoderms. These phyletic groups shared a common ancestor approximately 560 Mya, and while CNEs in these phyla show no conservation between them, they are highly predictive of the extent of the regulatory domains of homologous genes in both lineages. This association is observable both at the level of individual loci and genome-wide, independent of how the topological organisation of the genome is represented.

## TADs associated with GRBs exhibit distinct genomic features

Genomic regions have previously been classified into TADs, inter-TADs and TAD boundaries based on their size and interaction structure [26]. Several genomic features potentially involved in delimiting TAD boundaries have been identified including gene density, CTCF binding and the distribution of repetitive elements. We have previously reported that GRBs do not cover the whole genome and that many genes fall outside of regions with high-levels of non-coding conservation [17, 20], suggesting differences in the structural and epigenetic organisation of these regions. For this reason, we defined the set of TADs associated with high levels of extreme non-coding conservation as GRB-TADs, and attempted to identify features which distinguish them from TADs lacking evidence of this type of conservation (nonGRB-TADs) (see Methods).

Previously, it has been found that the regions surrounding key developmental genes are depleted of transposons [56], suggesting that the regulation of these genes is highly sensitive to insertions. Indeed, GRBs sharply define regions depleted of transposons, with strong increases in the density of SINEs in their flanking regions (Fig. 4, S7A). Changes in SINE density have previously been found to be associated with TAD boundaries [26]. It would appear that the regions associated with GRBs are associated with lower levels of retrotransposons compared to all TADs and nonGRB-TADs (Fig. 4A), reflecting selective pressure on both the syntenic and regulatory conformation of these loci. This supports SINE density as a marker of TAD boundaries, but primarily at GRB-TADs, and suggests that this pattern is the result of the selective pressure against transposon insertion disrupting the regulatory landscape of TADs containing developmental regulators.

Filion *et al.* used DamID to map multiple DNA binding proteins and histone modifications in *Drosophila*, classifying chromatin into five putative states/colours (see Methods) [57]. Using this classification, dm3-droMoj3 GRBs show the presence of transcriptionally silent regions (black), Polycomb-repressed chromatin (blue), or regulated euchromatin (red) within them (Fig. 4B, S7B), along with a clear depletion of constitutive heterochromatin (green). Therefore, the majority of GRBs in Kc167 cells are either silent or largely repressed by Polycomb. Indeed the domains of black and blue chromatin closely correspond to the extent of GRBs, with the edges of GRBs showing evidence of enrichment of constitutively active (yellow) chromatin. This enrichment for yellow chromatin is analogous to the presence of ubiquitously expressed/housekeeping genes at TAD boundaries as reported previously [26]. The pattern of regulated chromatin at GRBs is concordant with the expected expression pattern of GRB target genes, which are repressed by Polycomb in the majority of tissues and marked with red chromatin when active. This suggests that GRB-TADs are coherent with respect to their chromatin state and correspond to demarcated regions of regulated chromatin, which are often flanked by domains of constitutively active chromatin.

**Figure 3:**
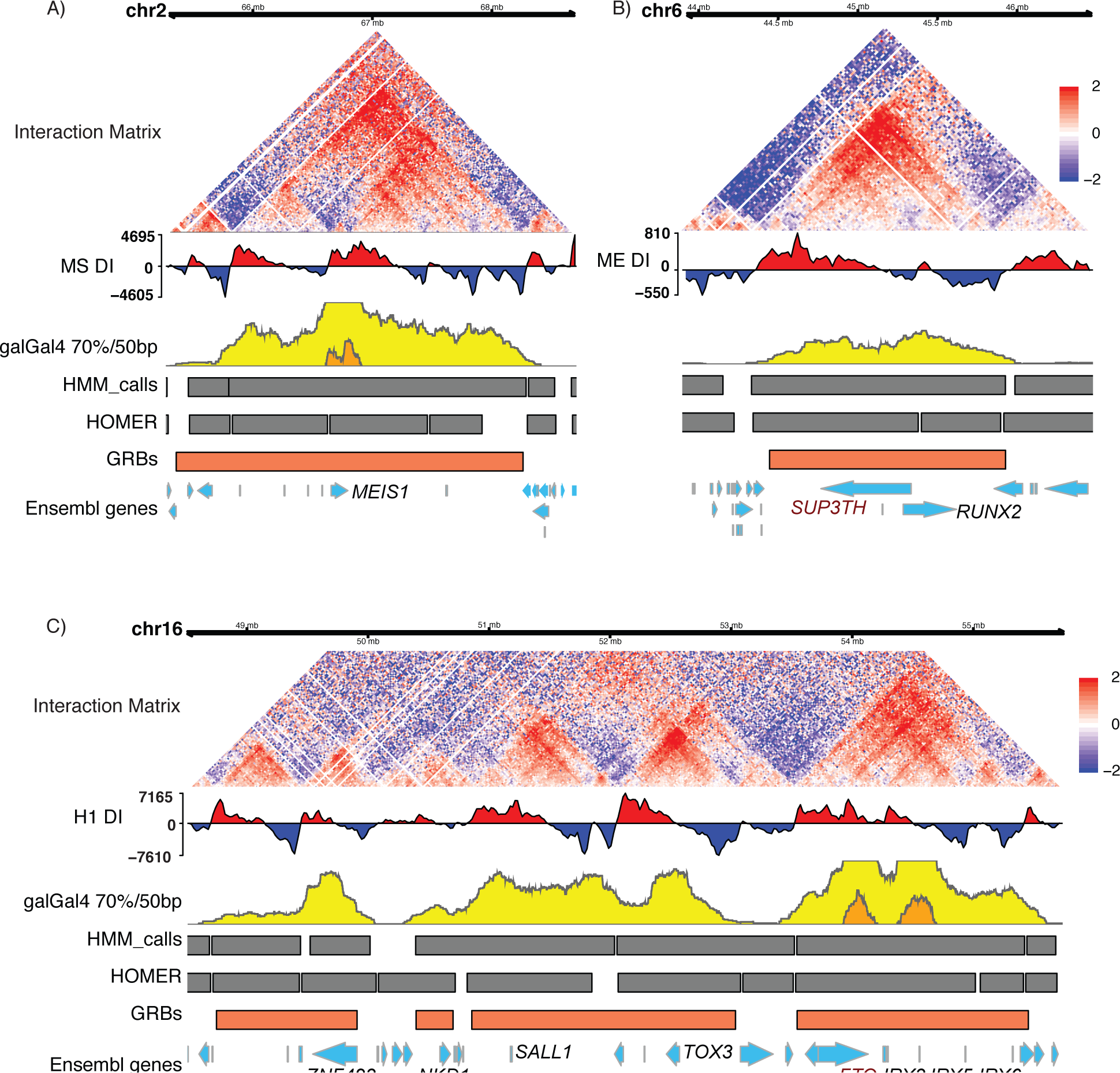
Examples of genomic regulatory blocks and their associated interaction landscapes in human. GRBs at several human loci show strong association with the structure of regulatory domains proposed from Hi-C. A) The GRB containing *MEIS1* (chr2:65270920-68723490), accurately predicts the span of regulatory interactions defined by Hi-C. B) The region located at chr6-44198640-46071520 contains both the transcription factor *RUNX2* and its bystander gene *SUP3TH* (shown in brown), both of which are located within a GRB which predicts the topological organisation of the locus. C) A region located on chr16:48476700-55776880 in hg19 contains several GRBs containing important developmental regulators, including *IRX3/5/6*, *TOX3*, *SALL1*, *NKD1* and *ZNF423*, which exhibit strong concordance with TADs. The *IRX3/5/6* locus contains homeobox proteins which have multiple functions during animal development and contains a well known bystander gene *FTO* (shown in brown), which contains an intronic enhancer which drives expression of *IRX3* [14, 50, 51, 55].

**Figure 4:**
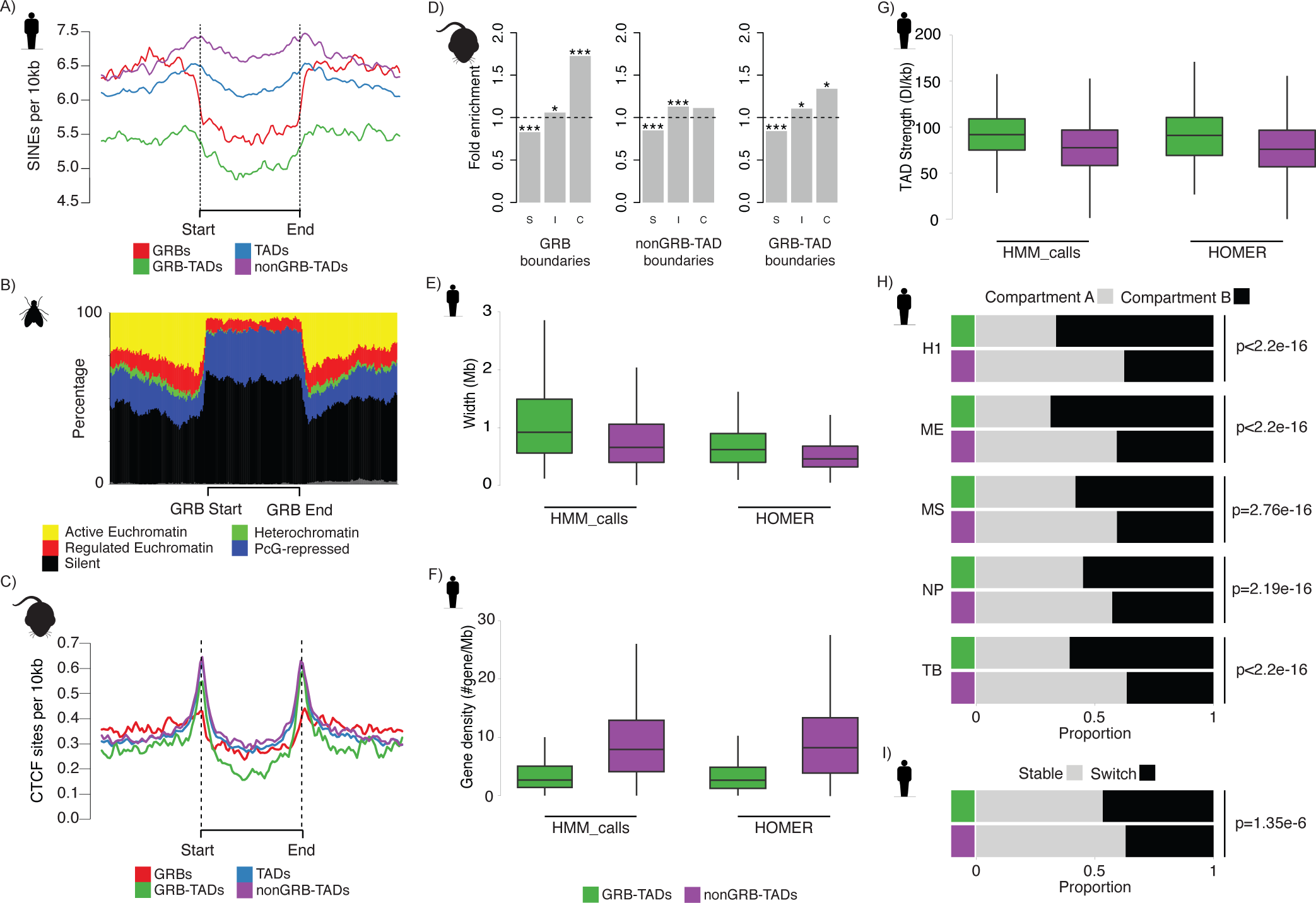
Several sets of features distinguish between TADs associated with extreme conservation (GRB-TADs) from those without (nonGRB-TADs) A) Depletion of SINE elements within GRB-TADs compared to nonGRB-TADs (using H1 HOMER TADs), reflects the selective constraint on these regions against repeat element insertion. B) GRBs in *Drosophila* Kc167 cells are mainly associated with inactive (black) and Polycomb-repressed chromatin (blue) and represent functionally coherent regions. Active and regulated chromatin correspond to different types of euchromatin. There appears to be a change in the proportion of constitutively active chromatin (yellow) at the boundaries regions identified as GRBs. C) CTCF sites are depleted within GRBs. CTCF sites per 10kb plotted across GRBs and flanking regions of equivalent size to the GRBs, normalised to show signal relative to GRB boundaries and loess-smoothed. D) Enrichment for different patterns of CTCF binding at different genomic features (Specific, Intermediate, Constitutive). Constitutive CTCF peaks are enriched within 10kb of both GRBs and GRB-TAD boundaries. (Binomial test p values: * p < 0.05, ** p < 0.01, *** p < 0.001). E) Distribution of TAD width reveals GRB-TADs are significantly longer than nonGRB-TADs identified in human H1 cells using either HOMER (median width 620kb vs. 460kbp, p <1e-6) or HMM calls (median width 920kb vs. 680kb, p <1e-6). F) Human H1 GRB-TADs are associated with lower protein-coding gene density than nonGRB-TADs identified using HOMER (median # genes 2.63 vs. 8.33 p <1e-6) or HMM calls (median # genes 2.65 vs. 8.33 p <1e-6). G) GRB-TADs are significantly stronger than nonGRB-TADs identified in human H1 cells using HOMER (median strength 90.88 vs. 75.81, p <1e-6) or HMM calls (median strength 91.77 vs. 77.68, p <1e-6). H) GRB-TADs (HOMER) are preferentially associated with Compartment B in all of the lineages investigated. I) GRB-TADs are more likely to switch compartment in at least one of the five lineages investigated (i.e. A-B or B-A) than nonGRB-TADs.

Next, we examined the relationship between the boundaries of GRB-TADs and the patterns of CTCF binding across multiple mouse tissues at different stages of development [58]. CTCF binding is depleted inside GRBs identified between mouse and chicken (Fig. 4C), however there is enrichment of CTCF peaks at the boundaries of GRBs, similar to the previously observed enrichment at TAD edges (Fig. S7C) [26]. This enrichment is more prominent at the edges of TADs that overlap GRBs than at the edges of the GRBs themselves, as expected if the predicted GRB boundaries tend to fall inside a TAD. Importantly, CTCF peaks at GRB boundaries and TAD boundaries are enriched for constitutive CTCF binding sites (p=4e-11, binomial test) and depleted for cell-type-specific CTCF binding (p=1e-8, binomial test) (Fig. 4D), in agreement with the invariance of TAD structure across cell types. CTCF peaks at GRB-TAD boundaries are enriched for constitutive CTCF binding (p=0.02, binomial test), while nonGRB-TADs show no significant enrichment for constitutive binding (p=0.11, binomial test), suggesting that the boundaries of GRB-TADs may be more consistent across different cell lines and tissues. This supports CTCF as being involved in the organisation and demarcation of these TADs, and the importance of insulating these domains containing key developmental regulators from interactions with non-cognate regulatory elements [59, 60, 61, 54].

In addition, GRB-TADs are significantly longer (p <1e-6, permutation test) than nonGRB-TADs in both human (Fig. 4E) and *Drosophila* (Fig. S7D), regardless of the method used to identify TADs. Despite their larger size, GRB-TADs tend to contain a lower number of genes than nonGRB-TADs (Fig. 4G, Fig. S7E). This is expected, as GRBs are associated with gene deserts [62]. The combination of these features suggests that these regulatory domains are larger to accommodate the numerous regulatory elements needed to spatiotemporally regulate target genes during development. In addition, examination of their directionality indexes showed that GRB-TADs are amongst the strongest in the genome in human (p <1e-6, permutation test, Fig. 4F) and *Drosophila* (p <1e-6, permutation test, Fig. S7F). This suggests that GRB-TADs have higher levels of self-interaction and are more insulated from neighbouring regions compared to nonGRB-TADs.

Since GRB-TADs are gene-sparse and their target genes are inactive in most tissues, we expect most GRBs to be associated with compartment B [25]. Indeed, regardless of cell lineage, GRB-TADs were found to be preferentially located within the B compartment of the nucleus (Fig. 4H, Fig. S8A). Dixon *et al.* reported that TADs that change compartment show concordant changes in expression of their constituent genes during development/differentiation [31]. We found a clear enrichment of GRB-TADs in the set of TADs that switch compartment in one or more of the cell types investigated compared to nonGRB-TADs which appeared to be more stable (p=1.35e-06, Fishers exact test, Fig. 4I, Fig. S8B). At several GRBs, a change in compartment was associated with the change in expression state of the GRB target gene (e.g. *ZEB2*, *OTX2* and *SOX2* Fig. S8D/E/F). Intriguingly, while the target gene showing concordant changes between expression status and compartment, nearby bystander genes showed no evidence of this relationship. While *ZEB2* (Fig. S8D) showed high expression in mesenchymal stem cells, in line with its presence in compartment A, other genes within this GRB exhibited little or no change in their expression. This was also apparent at the *OTX2* GRB (Fig. S8E) in mesendoderm and neural progenitor cells, and at the *SOX2* GRB in neural progenitor cells (Fig. S8F). This relationship between GRB-TADs and compartments further confirms that these regions represent the regulatory domains of developmental genes.

Several TADs not associated with GRBs identified using human-chicken comparisons nevertheless have features associated with GRB-TADs and contain known developmental regulators. At a number of loci, human-chicken CNEs are present but were not effectively clustered by our approach, however several loci appear to lack CNEs identifiable between human and chicken. The *CNTNAP4* locus (Fig. S9A) is largely devoid of CNEs identified using chicken, but shows limited conservation between human and spotted gar and high levels of non-coding conservation in comparisons involving dog (canFam3) and mouse. It appears that this regulatory domain is conserved over vertebrates, with the evolutionary patterns of its constitutive CNEs suggesting that these elements have been lost and gained in a lineage-specific fashion [10]. Investigating those nonGRB-TADs that switched compartment during differentiation identified several loci that exhibited high levels of non-coding conservation in comparisons with more closely related species (e.g. *NECTIN3* and *protocadherin alpha/beta* clusters). Enrichment analysis of this set of genes revealed a strong enrichment for genes involved in cell:cell adhesion (Fig. S8C). This enrichment suggests that these regions may represent loci which have undergone more recent regulatory innovation [63], or whose regulatory elements are subject to high turnover [10].

These results support the idea that TADs associated with developmental genes and high levels of non-coding conservation have a distinct set of features indicating differences in the topological organisation and demarcation of these regulatory domains. We conclude that most TADs called in CNE-free and gene-dense regions of the genome exhibit reduced directionality of interactions compared to those associated with high levels of non-coding conservation, which represents a distinct form of regulatory domain.

## The sizes of GRBs and TADs scale with genome size

Next we investigated whether there was a relationship between genome size, TAD size and GRB size. Previously it has been shown that clusters of CNEs are on average much more compact in *Drosophila* than in mammals [3], and that they scale with genome size in fish [64]. As expected, TADs and GRBs were larger in species with larger genomes (Fig. 5A). Therefore, we investigated whether the genomic locations of CNEs could provide information on the expansion and evolution of TADs in vertebrates.

We investigated a stringent set of seventeen human GRBs which accurately predicted the boundaries (i.e. both edges within 120kb) of the same set of TADs regardless of which species was used to identify GRBs. These include the GRBs containing *MEIS1* (Fig1, Fig, 3), IRX3 (Fig. S2A, Fig. 3), and other important developmental genes such as *PBX1*, *OTX1* and *LMO4*. The location of the homologous CNEs was used to predict an orthologous set of GRBs in spotted gar, chicken and opossum. In spotted gar this set of GRBs was 24% of the sizes of GRBs predicted in human using the same set, with the ratio of genome sizes being 28%. This relationship was also observed in chicken (median GRB size ratio 37%, genome size ratio 33%) and opossum (median GRB size ratio 103%, genome size ratio 113%). Indeed, overall the size of these regions in one species was highly predictive of the size of the region in another (Fig. 5C). The ratios of these numbers and their strong linear relationship suggest that this set of domains has undergone expansion at a comparable rate to genome growth. This relationship suggests that the distribution of CNEs in one species could be used to predict the location of GRBs and their associated TADs in another.

However, this pattern of expansion and evolution is not the only one observed in our analysis: some regions exhibit strong conservation of one GRB boundary whilst showing expansion at the other boundary, e.g. *HLX* locus (Fig. S9B), while others have a core highly conserved region in comparisons with spotted gar which is expanded in dog and opossum comparisons, e.g. *CNTNAP4* (Fig. S9A). These observations support the idea of multiple mechanisms involved in the evolution of regulatory domains in Metazoa including lineagespecific expansion of TADs, recruitment of neighbouring TADs and recruitment or turnover of regulatory elements both within these domains and at their edges. Future Hi-C datasets from evolutionarily distant species will enable a detailed investigation of the relationship between the evolutionary patterns of non-coding conservation and topological organisation.

## Discussion

In this work, we have presented multiple lines of evidence for the equivalence of a functionally distinct subset of TADs with regions of long-range regulation defined by clusters of extremely conserved noncoding elements (known as genomic regulatory blocks, or GRBs). We show that the span of clusters of CNEs is predictive of the span of a subset of TADs in both human and *Drosophila*. This set of TADs, referred to as GRB-TADs, show a set of features that appears to distinguish them from TADs lacking CNEs and apparent long-range regulation. These GRB-TADs show distinct patterns of retrotransposon density and CTCF binding, and are significantly longer and associated with denser interactions than TADs lacking CNEs. We conclude that this set of TADs are evolutionarily ancient 3D structures predicted by clusters of extreme non-coding conservation, which contain genes involved in the regulation of embryonic development and morphogenesis and their associated regulatory elements. This relationship presents genome-wide evidence for TADs as structural and regulatory domains around key developmental genes under long-range regulation. The connection between the distribution of CNEs and topological organisation has far-reaching implications for understanding the nature and evolution of long-range regulation at developmental loci.

The specific set of features associated with those TADs exhibiting high levels of extreme non-coding conservation, referred to as GRB-TADs, suggests that these correspond to a distinct functional class of regulatory domains. TADs associated with extreme non-coding conservation are some of the largest, strongest and most gene-sparse in human and *Drosophila*. This suggests that chromatin structure is mainly defined by the need to regulate important developmental genes, with nonGRB-TADs being smaller, weaker, and less important in determining overall chromatin structure. Since GRBs form the regulatory domains of developmental genes, in any one tissue or developmental stage most GRBs will be inactive and marked by Polycomb and H3K27me3 [65, 66]. The degree of intermixing of chromatin between genomic domains depends strongly on their epigenetic state, with Polycomb-repressed chromatin showing little or no spatial overlap with active or inactive chromatin [67]. It has also been suggested that regions of active chromatin within TADs interfere with the packaging of chromatin, disrupting TAD formation and leading to less compact TADs, or fragmentation into smaller TADs [68]. The enrichment of active euchromatin at the boundaries of these regions may reflect the formation of barriers. The lower density of active genes and increased likelihood of Polycomb repression at GRBs may explain some of their topological features.

**Figure 5:**
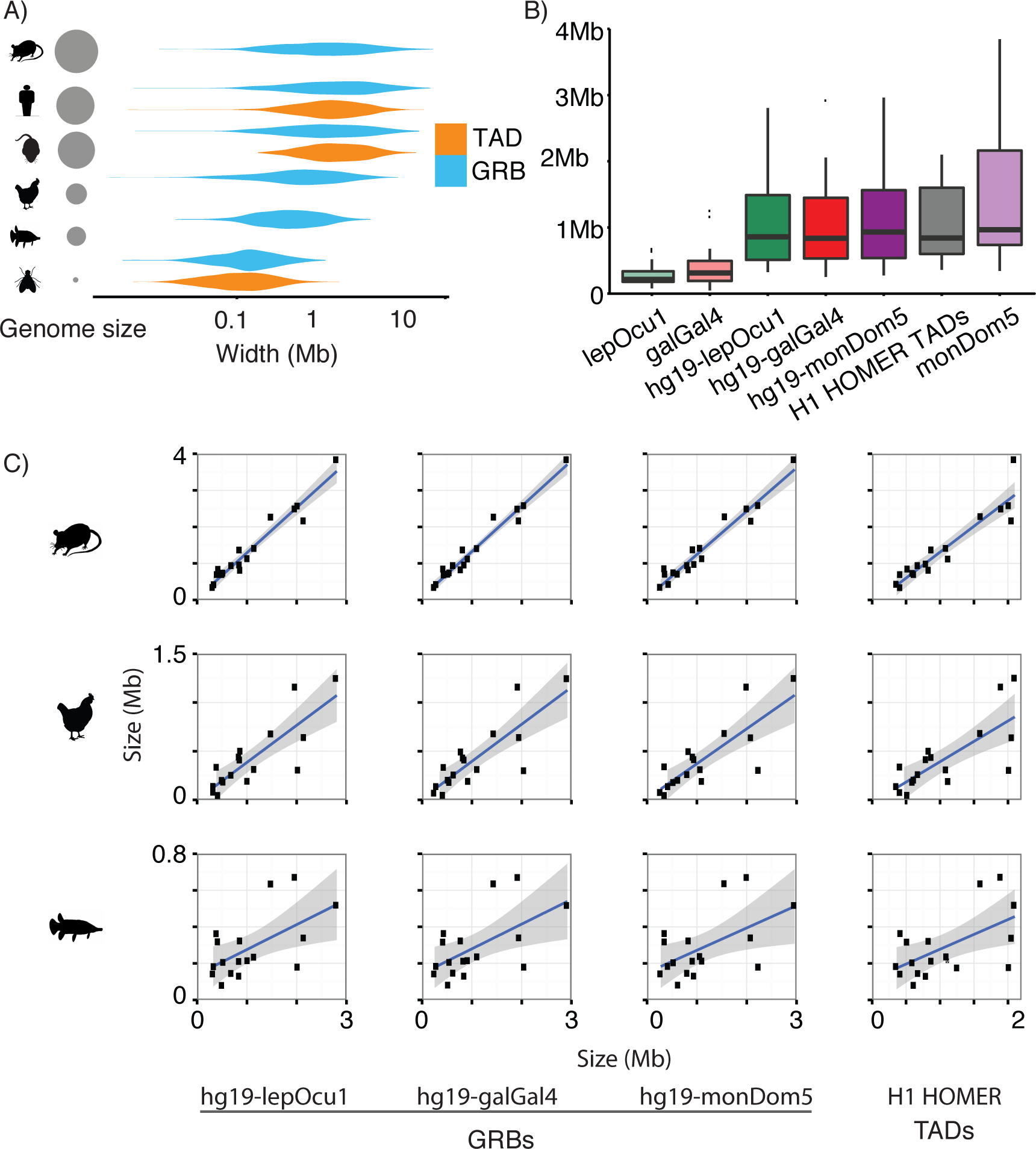
The span of CNEs in various species is predictive of genome size and TAD size. A) Distribution of genome size, TAD size and GRB size in opossum, human, mouse, chicken, spotted gar and *Drosophila*. B) Sizes of 17 stringent GRBs identified in various species, shows direct expansion and growth in line with that expected given differences in genome size C) Correlation of sizes of GRBs identified in one species compared to other species and to the corresponding TADs in human, finds that the size of a GRBs identified in other species is highly predictive of the size and scope of regulatory domains in other species.

The conservation and divergence of CTCF binding sites is thought to play important roles in the evolution of regulatory domains [54, 33]. Constitutive CTCF sites are enriched at GRB and GRB-TAD boundaries, suggesting these regions are consistently insulated from neighbouring domains. However, the overall role of CTCF is unclear, as there is additional CTCF binding within a GRB/GRB-TAD, although at lower levels compared to other TADs. This may reflect its ability to function as both an insulator and organiser of chromatin interactions [69, 70]. The enrichment of CTCF at GRB and GRB-TAD boundaries, combined with the strength of interactions associated with GRBs, suggests that these regions do not strongly interact with elements in adjacent regulatory domains. This is in addition to the association of GRBs with Polycomb-repressed chromatin colours and depletion of active euchromatin, which also promote insulation from neighbouring regions, as described above. These features are reflective of the multiple mechanisms involved in insulating target genes from ectopic regulation by elements in neighbouring domains.

GRBs and TADs are both stably present across different cell types, however TADs are also organised into inactive and active chromatin compartments, and the assignment of TADs to compartments can vary across cell types [25, 26, 71, 27, 31]. GRB-TADs are preferentially located within the inactive B compartment across all five of the lineages investigated. This is expected given that GRBs contain developmentally regulated genes, whose expression is constrained to specific cell lineages. This indicates that TADs which are associated with compartment B have increased non-coding conservation compared to TADs which are stably present in the A compartment. We also observed that GRB-TADs are more likely to switch compartments than those TADs lacking extreme non-coding conservation, and we identified changes in the expression of genes within GRBs that are concordant with their assignment to the active or inactive compartment (Fig. S8D-F). Importantly, these changes in gene expression are in several cases restricted to the GRB target gene, while other genes within the GRB are unaffected by the compartment switch. This is consistent with our hypothesis that GRBs represent regions that encompass regulatory elements of specific developmental regulators, which are repressed during the majority of developmental stages and cell types and only expressed in a limited subset.

It has recently been reported that the 3D structure of developmentally regulated genes in *Drosophila* is pre-configured, with linearly distant enhancers found in close spatial proximity to their target gene even before the activation of enhancers and target gene expression [72]. This spatial preconfiguration is apparently a phenomenon distinct from the dynamic chromatin contacts detected by FISH [73]. This raises the possibility that interactions within TADs corresponding to GRBs could be pre-configured to constrain regulatory elements or channel their interactions in all cells, regardless of the expression status of the target gene, and are therefore more stable across cell types than interactions within other TADs.

The striking concordance between strong TADs, which are invariant across species and cell types, and GRBs, provides multiple lines of evidence that at these loci they reflect the same underlying functional and evolutionary phenomenon. Combining the concepts of GRBs and TADs leads to a model of regulatory domains with stronger predictive and explanatory power than either concept alone (Fig. S10). The GRB model provides a framework for long-range regulation, in which the majority of regulatory elements within a GRB are dedicated to the control of its target gene [16, 13], with other genes not responding to long-range regulation despite being located close to regulatory elements [12, 74]. Target and bystander genes appear to exhibit distinct features that may help to explain this specificity [20, 21]. Topological organisation data provides more precise boundary estimates for these regions, including the ability to separate adjacent GRBs, and contributes information about the stability of the organisation of these domains and regulatory interactions within them across cells and tissue types. Therefore, a TAD enriched for extreme non-coding conservation is not representative of the regulatory domain of all of its constitutive genes, but primarily corresponds to the regulatory domain of the target gene under long-range regulation.

Our estimated spans of GRBs are of limited precision and coverage, but we have shown that the distribution of CNEs can serve as a proxy for the extent of a functionally distinct subset of TADs in both human and *Drosophila*. Therefore, clusters of CNEs identified between evolutionarily distant species could be used to infer regulatory domains and to predict the topological organisation of these domains in species lacking Hi-C data. At several loci, it appears that the boundaries of topological domains predicted using CNEs are highly similar regardless of the species involved in generating that prediction. This observation shows that not only are regulatory elements in these regions under intense selective pressure, but that the basic structure of these loci has existed over hundreds of millions of years of evolution.

The association between the boundaries of GRBs and TADs, the conservation of GRB boundaries and the structural features of the overlapping TADs suggests that 3D organisation at these regions is highly conserved and under intense selective pressure, at least back to the ancestor of tetrapods and arthropods. Regions containing homologous developmental genes are associated with the same type of conservation and structure in human and *Drosophila*. However, the phenomenon of microsynteny has existed since early Metazoa and is apparent across bilateria [23, 75, 76]. The syntenic relationship between *RUNX2* and *SUPT3H* (Fig. 3A) is conserved between humans and sponges [77], and a GRB containing Iroquois and Sowah genes is conserved across a wide range of bilaterians, apart from tetrapods [78]. This suggests that the presence of this distinct form of regulatory domain is ancient, and may have existed since the origin of Metazoa.

Experiments investigating chromatin conformation in eukaryotes other than Metazoa, such as the yeast Saccharomyces cerevisiae and the plant Arabidopsis thaliana, have reported that their genomes are not segmented into TADs [79, 40]. These organisms lack most or all of the developmental regulatory processes mediated by GRB target genes in Metazoa. Intriguingly, the Arabidopsis genome has TAD-like structures at some loci, which have been found to harbour genes marked by H3K27me3, suggesting their involvement in the regulation of plant development. Whether this means that the same structural phenomenon underpins the regulation of multicellular development remains to be investigated.

This relationship suggests that the regulatory and topological domains around developmental genes are dependent on conserved genome sequence, and therefore likely to be stable across evolution as well as across cell types. A remarkable property of the correspondence between TADs and GRBs is that they expand and shrink along with the entire genome, suggesting that the 3D organisation of regulatory loci is robust towards gain and loss of DNA between its constituent CNEs, even though insertion of repetitive elements is disfavoured. Previously, we observed that GRBs are on average much more compact in *Drosophila* than in mammals [3], and that they also scale with genome size in fish genomes [64]. The correspondence of GRBs and TADs predicts that TADs are smaller in *Drosophila*, as previously observed [38]. Duplicated loci containing developmental regulators are more likely to be retained after whole genome duplication (WGD) [80, 81]. Therefore, since tetrapod lineages have undergone two WGDs, a larger number of developmental regulators and associated GRBs are expected compared to arthropods, which is confirmed by our analysis.

The expansion and shrinkage of TADs and the turnover of regulatory elements within them is likely to be a highly important mechanism in metazoan evolution [82, 83] and be responsible for considerable differences in organismal complexity [84, 85, 86, 87, 10]. The syntenic organisation of *SHH* with LBMR1 and RNF32 is a vertebrate-specific evolutionary innovation which is required for proper regulation of *SHH* in the limb bud via an enhancer located within an intron of *LMBR1* [18, 88]. Experiments which have disrupted the boundaries of TADs have found that this severely alters the spatial organisation of the locus, allowing ectopic enhancer-promoter interactions and leading to aberrant expression of target genes [54, 30]. Deletions within TADs can lead to changes in enhancer-promoter interactions, in some cases causing disease [89, 90, 91], which suggests some level of selective pressure against such re-arrangements. The depletion of retrotransposons within GRB-TADs suggests that their insertion within this type of regulatory domain is under intense negative selection. This pressure may be due to the ability for retrotransposons to create new cis-regulatory elements [92], potentially perturbing the organisation of interactions within a regulatory domain resulting in a negative effect on fitness in the majority of cases [93]. In future, analysis of the evolutionary dynamics of CNEs and genomic rearrangements within TADs in multiple species will help to provide insights into their evolutionary dynamics.

The reported spatial correspondence of GRBs and TADs does not offer immediate suggestions for the elusive origin of extreme non-coding conservation. Current models of genome folding do not include a mechanism that could account for this level of selective pressure on elements within TADs. Since our results strengthen our ideas that GRBs are unified functional structures of long-range regulation, the next step will be to decipher how the sequence information contained within them is used to establish their stable 3D structure at the beginning of each cell cycle and how these structures preconfigure the regulatory environment of their target genes, allowing their precise regulation during development.

## Methods

### Identification of conserved non-coding elements

Conserved non-coding elements (CNEs) were generated by examining pairwise BLASTZ net whole-genome alignments [94] for regions with a high percentage identity over a defined number of base-pairs. For each comparison, both of the relevant nets (from the perspective of each species) were scanned. Elements overlapping exonic and repetitive repeats were removed. This set of elements was then aligned against the genome using BLAT to remove elements that mapped to more than four locations in vertebrates. The resulting set of CNEs was then smoothed using a sliding window (300kb for vertebrates and 50kb for *Drosophila*) to generate CNE densities, as was originally used for ANCORA browser [95].

### Identification of Genomic Regulatory Blocks (GRBs)

GRBs were generated by identifying regions of the genome that contain a high density of syntenic conserved non-coding elements (CNEs). CNE densities were generated using CNEs that showed a high level of percentage identity (i.e. at least 70-96%) over a number of base pairs (i.e. 30bp or 50bp) between two species. These densities were partitioned into regions with high or low CNE density using an unsupervised two-state HMM [96] (Fig. S1). This segmentation was performed ten times, with the set having the best Akaike information criterion (AIC) defined as the best model. All CNEs that were not present in an enriched region were removed. The remaining CNEs were merged using the distance between adjacent CNEs as a criterion. The sizes of gaps between individual CNEs for each chromosome were determined and used to recursively split the genome into regions where the distance between adjacent CNEs was greater than a specified quantile of the gap distribution. Our experiments with this threshold suggested that as the evolutionary distance between the two species of interest increased, the quantile used needed to be decreased. We used 0.98 for hg19-monDom5 comparisons, 0.98 for hg19-galGal4, and 0.93 for hg19-lepOcu1 and 0.97 for dm3-droMoj3. These parameters were determined empirically by investigating the ability of our predictions to recapitulate known boundaries of GRBs. Following this, putative regions were split by the chromosome that they originated from in the query species to generate discrete regions of conserved synteny. Regions that did not contain a protein-coding gene were merged with adjacent regions if they were within 300kb (for human/mouse) or 50kb (for *Drosophila*). All remaining regions not containing a protein-coding gene or having fewer than ten CNEs was discarded, resulting in a set of putative GRBs.

Our initial investigations found that segmenting CNE density using either a HMM or the distance-based criterion alone had difficulties generating a robust set of putative GRBs. The HMM approach had a tendency to merge adjacent GRBs, while the distance-based clustering appeared to be very sensitive to isolated CNEs and tended to overestimate the span of GRBs. By combining both methods together, the impact of the problems associated with each method was reduced.

### Processing of Hi-C datasets

The set of TADs generated from an experiment is dependent on both the experimental protocol (i.e. restriction enzyme, sequencing depth) and on the processing techniques used (i.e. bin size, algorithm). Hi-C interaction datasets for human were obtained from GEO (GSE52457), for H1-ESC (H1), mesenchymal stem cells (MS), mesendoderm (ME), neural progenitor cells (NP) and trophoblast-like (TB) [31]. These reads were iteratively aligned [97] using bowtie [98] against hg19. Reads mapping to chrM and chrY were removed from the analysis. The resulting aligned reads were binned using a variety of bin and window sizes, with a bin size of 20kb and a window size of 40kb appearing to generate a robust set of TADs. TADs were identified using both HOMER [42] and the TAD calling pipeline (HMM calls) proposed by Dixon *et al.* [26]. Mouse ESC Hi-C data was downloaded from GEO (GSE35156) and processed using the same pipeline as for human. Hi-C for *Drosophila* whole embryo Hi-C data [38] was obtained from GEO (GSM849422) and processed using the same pipeline. Directionality matrices for *Drosophila* Hi-C were generated using HOMER with a bin size of 10kb and a window size of 20kb. Reads mapping to heterochromatic chromosomes (i.e. chr3RHet, chr2LHet) were discarded from further analyses. TADs predicted to span across centromeric regions were removed.

The strength of TADs was defined as the sum of the absolute directionality indexes within a TAD normalised to the length of the TAD in kilobases.

High-resolution GM12878 Hi-C data was obtained from GSE63525 [27]. Contact domains which overlapped by at least 60% were collapsed to generate a set of outer-most domains.

Compartments were identified by performing PCA on the Hi-C interaction matrix and investigating the 1st principal component. TADs were classified as A or B given at least 60% of locations within them were either positive or negative, respectively. A single gene was classified by examining at 5kb window around its promoter and classifying it as belonging to the A or B compartment using the same criteria.

### Calculating expected number of GRB boundaries lying within a specific distance of TAD boundaries

The significance of the number of GRBs having both edges within three Hi-C bins of a TAD boundary was calculated by comparing the observed number of GRBs with that observed by randomly shuffling GRBs. A null distribution was created by generating 1000 shuffled regions using BEDtools [99], excluding centromeric regions, and calculating the number of random regions whose boundaries were within three Hi-C bins of the nearest TAD boundary.

### H3K27ac data

H3K27ac data for human H1, ME, MS, NP and TB cells was obtained from GSE16256 [100], and aligned against hg19 using bowtie [98]. Enriched regions were identified using MACS2 [101], using default parameters. ChIP dataset quality was assessed using ChIPQC [102], and the dataset with the highest ChIP enrichment was used (identified as the replicate having the maximum RiP (reads in peaks)). Duplicated reads were removed from each dataset, and the resulting coverage was then normalised for read depth. The average coverage within a 5kb bins was then calculated around putative GRBs.

### GRB-TADs vs nonGRB-TADs

TADs were classified as GRB-TADs and nonGRB-TADs based on their overlap with GRBs. In human, TADs with more than 80% overlap with a single GRB were assigned as GRB-TADs, TADs with less than 20% overlap with a TAD were assigned as nonGRB-TADs, and any TAD that overlapped more than one GRB or had an overlap percentage between 20% and 80% was screened out of this analysis. For *Drosophila*, a TAD with greater than 60% overlap with a single GRB was designated as a GRB-TAD, with TADs showing less than 25% overlap with a GRB defined as a nonGRB-TAD. These thresholds were determined by investigating the distribution of percentage overlap between GRBs and TADs genome-wide.

The significance of the differences between median TAD width, gene density and strength were calculated empirically by randomly permuting the labels of TADs 1 million permutations to generate a null distribution for each statistic.

### RNA-seq analysis

RNA-seq data was obtained from GSE16256 [103], and aligned against Ensembl genes release 75 using Tophat2 [104]. Aligned reads were counted using htseq [105]. Differential expression analysis was performed using DESeq2 [106].

### Repetitive elements

Repetitive elements were obtained from the UCSC Genome Browser [107] for hg19 and dm3, with only those elements classified as SINEs used in the analysis. For visualisation using heatmaps the average coverage of SINE elements within 5kb bins in a 8Mb window around the centre of hg19-galGal4 GRBs was calculated. For average profiles, SINEs per 10kb were calculated by counting the number of SINE elements occurring in overlapping 10kb windows, with a step size of 1kb, across the human genome.

### CTCF binding at GRBs and TADs

For this analysis, we used GRBs called using mm9-galGal4 CNEs at a threshold of 70% over 50bp. CTCF ChIP-seq data was obtained from mouse ENCODE [58] for 17 cell lines and tissues. Reads were aligned to the mm9 genome using bowtie [98] and peaks called using MACS2 [101], with the first input replicate for each sample used as the control. Where replicates were available, the intersection of peaks called on different replicates was used for the final peak set.

A consensus set of CTCF peaks was calculated by resizing all peaks to a width of 400bp and taking the union of peaks across all 17 samples. Peaks were scored for the number of samples they occur in. CTCF peaks per 10kb tracks were calculated using the consensus peak set and counting the number of peaks occurring in overlapping 10kb windows, with a step size of 1kb, across the mouse genome.

CTCF peaks within 10kb of GRB and TAD boundaries were assigned to the boundary, and classified as ‘specific’ if they were present in 1-2 samples, ‘constitutive’ if they were present in 16-17 samples, and ‘intermediate’ otherwise. Enrichment was calculated relative to the proportion of these categories in the consensus peak set, and p-values calculated for each category using a two-sided binomial test.

### D. melanogaster chromatin states

*Drosophila* chromatin state data was downloaded from GEO (GSE22069). Filion *et al.* identified five chromatin states in *Drosophila* melanogaster Kc167 cells [57]. Each of these states was found to have specific characteristics, including transcriptional activity, replication timing and biochemical properties. With black chromatin associated with transcriptionally silent developmentally regulated chromatin, green with classical heterochromatin, and blue with chromatin bound by PcG proteins. Red chromatin was associated with early replicating euchromatin, a high density of regulatory elements and the presence of genes under complex, long-distance regulation. Whereas, yellow chromatin was associated with late-replicating euchromatin and genes with ubiquitous expression patterns (house-keeping genes). Heatmaps were generated by dividing a 1Mb region around the centre of dm3-droMoj3 GRBs into 1 kb bins, with each bin assigned the most common chromatin colour within it.

### Comparison of the relationship between genome, TAD and GRB size

A set of seventeen human GRBs that accurately predicted the boundaries (i.e. both edges within 120kb) of the same set of TADs regardless of the species involved (i.e. using monDom5, galGal4, lepOcu1) was identified. For each GRB, the location of the constitutive CNEs in the reference species was identified and used to generate a putative GRB in that species. Any CNEs that were found to be present within the GRB but originated from a separate part of a chromosome was removed.

### Data availability

A trackhub containing all of the data presented in this analysis is available at http://trackhub.genereg.net/harmston2016/harmston2016.hub.txt

## Acknowledgements

We thank Piotr Balwierz, Anja Barešić, Sarah Langley, Dimitris Polychronopoulos and Owen Rackham for their comments on the manuscript. N.H., E.I-S. and B.L. are supported by the Medical Research Council UK. G.T is supported by the Wellcome Trust. G.T. and B.L. are supported by EU project ZF-Health (FP7/2010-2015 grant agreement no 242048). MP by a MRC DTP supported by the Faculty of Medicine, Imperial College London.

## 1 Supplementary Figures

**Supplementary Figure 1:**
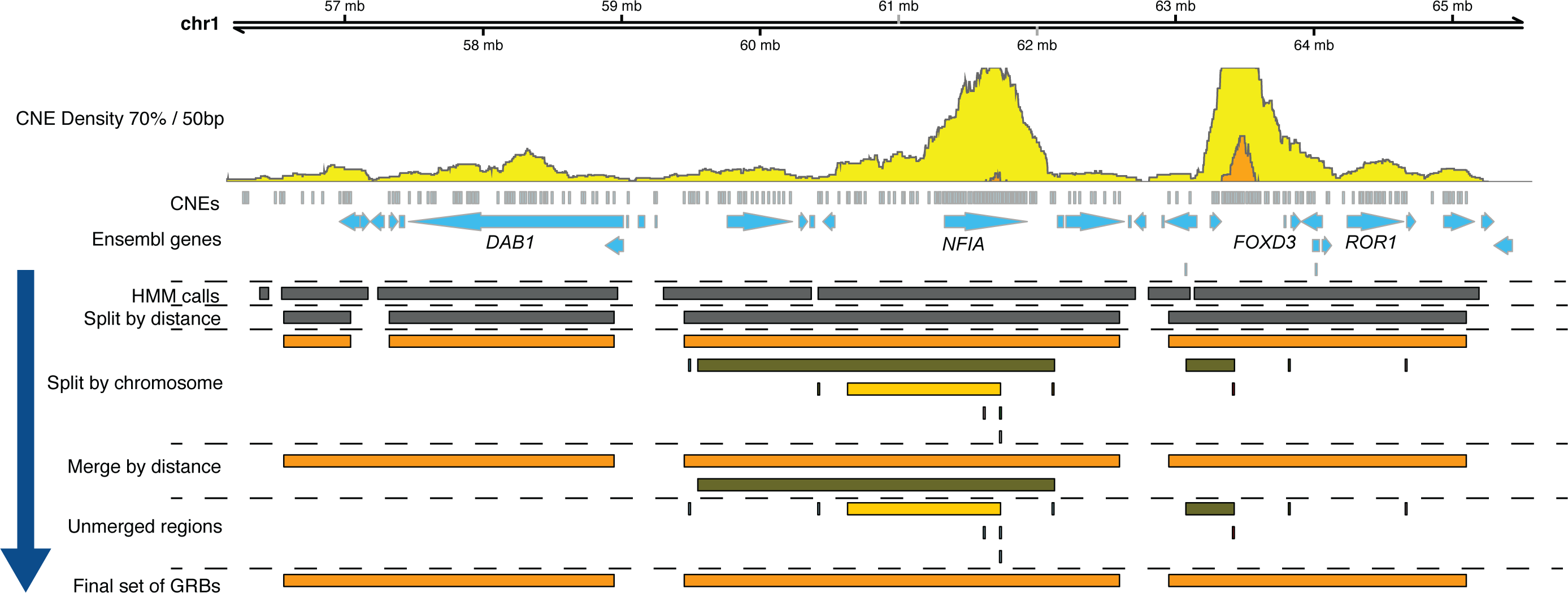
Schematic of the method for segmenting the genome into GRBs. The region chr1:56136940-65569360 contains several distinct regions of CNE density overlapping with known developmental regulators (*DAB1, NFIA, FOXD3*). Our CNE clustering method (*see* Methods) splits the CNE density obtained from ANCORA into distinct regions by first segmenting the density using a HMM into regions of high and low density, and then clustering the CNEs together using the distance between them. This is followed by the application of various heuristics to generate a set of putative GRBs. These regions are split by the chromosome that they originate from in the query species, and merged with nearby regions if they are from the same chromosome and within a specified distance. While we expect a number of GRBs to be incorrectly segmented, the resulting set of GRBs appear to match, via visual inspection, the known GRB architecture at a number of loci.

**Supplementary Figure 2:**
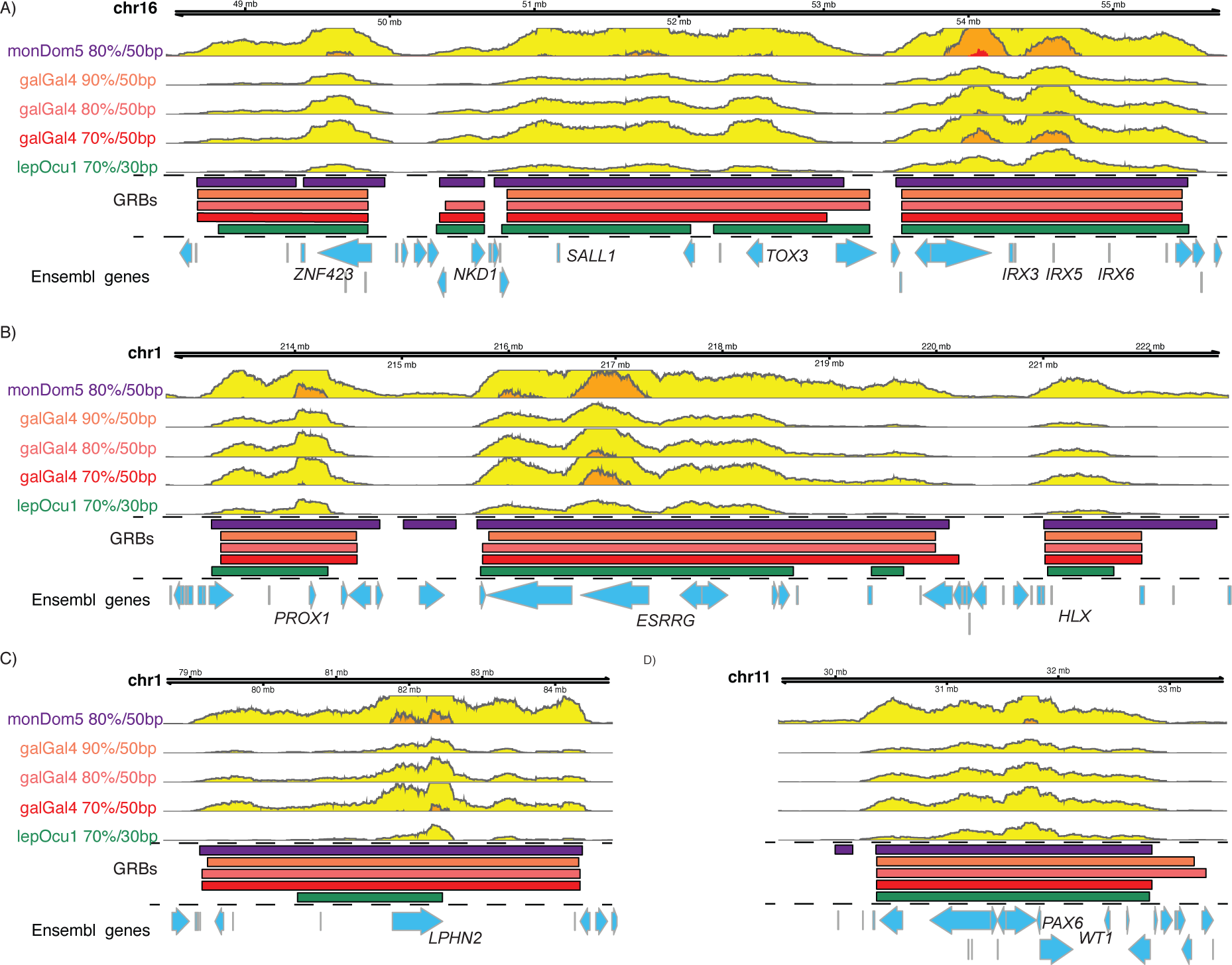
The boundaries of GRBs are highly consistent regardless of thresholds or species. A number of loci in human are enriched for CNEs identifiable between evolutionarily distant species at various thresholds. For each set of CNEs, the corresponding CNE density is displayed as a horizon plot along with the corresponding set of putative GRBs for each comparison. A) A large chromosomal region in human (chr16: 48476700-55776880) contains several GRBs which are highly conserved over multiple evolutionary comparisons. The edges of the predicted *IRX3/5/6* GRB are highly concordant between all species and thresholds investigated, with the other GRBs in this region showing strong agreement in the majority of cases. B) A region (chr1:212820540-222701560) containing three known developmental regulators (*PROX1*,*ESRRG* and *HLX*) is segmented into three GRBs. In the majority of cases at this region at least one boundary of a GRB appears to be identified consistently over all comparisons. C) The GRB (chr1: 78652340-84766720) around *LPHN2*, a latrophilin involved in cell adhesion, has similar boundaries in comparisons generated using monDom5 and galGal4 whilst the GRB identified using lepOcu1 is much smaller. D) While it is known that the region (chr11:29500000-33500000) containing *PAX6* and *WT1* forms two separate GRBs Navratilova:2009ip, we are unable to distinguish between these regions based on CNE density alone. One edge of this GRB appears to be robustly identifiable in all comparisons with the other edge exhibiting some variation.

**Supplementary Figure 3:**
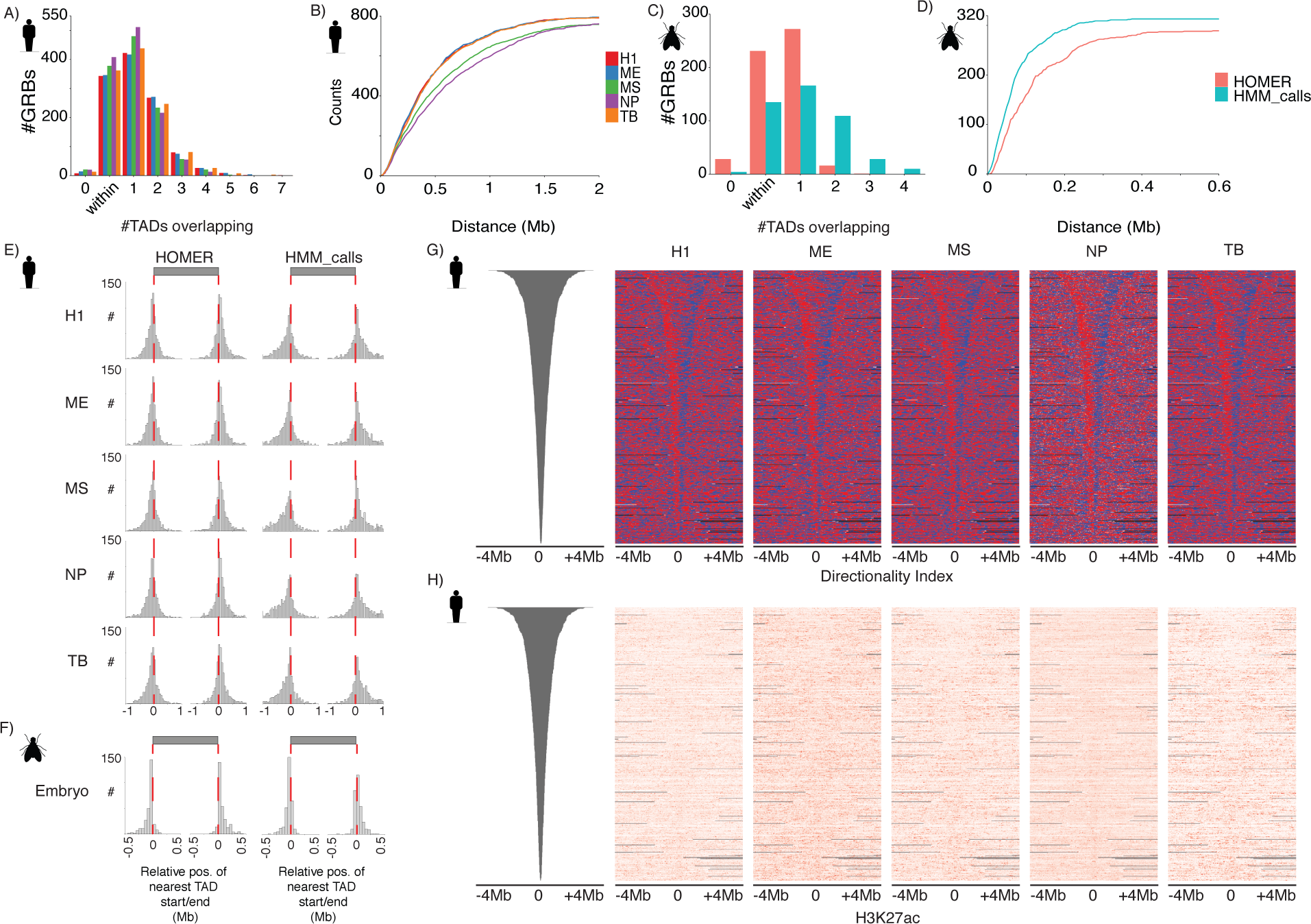
The boundaries of GRBs predict the boundaries of TADs identified in multiple cell lineages. A) A large number of hg19-galGal4 GRBs was found to be located within individual TADs (identified using HMM calls) or overlapping only a single TAD, regardless of cell lineage. B) Cumulative distribution of distance to nearest TAD (HMM calls) boundaries from GRB boundaries in different cell lineages considering both edges i.e. both the start and end position of a GRB lie within Xkb of the nearest TAD start and end. C). A large number of dm3-droMoj3 GRBs was found to be located within individual TADs (identified using HOMER and HMM calls) or overlapping only a single TAD. D) Cumulative distribution of distance to nearest TAD (HOMER and HMM calls) boundaries from GRB boundaries in *Drosophila* whole embryos considering both edges, i.e. both the start and end position of a GRB lie within Xkb of the nearest TAD start and end. E) Relative position of the nearest TAD start/end compared to the boundaries of hg19-galGal4 GRBs, using TADs identified in multiple cell lineages (H1-ESC (H1), mesenchymal stem cells (MS), mesendoderm (ME), neural progenitor cells (NP) and trophoblast-like (TB)). F) Relative position of the nearest TAD start/end compared to the boundaries of dm3-droMoj3 GRB, using TADs identified in *Drosophila* Hi-C data. G) Heatmaps representing overall direction of the Hi-C directionality index calculated in different cell lineages, spanning an 8Mb window around the centre of putative hg19-galGal4 GRBs. H) The distribution of H3K27ac signal in an 8Mb around the centre of hg19-galGal4 GRBs reveals no obvious strong association between these marks and the boundaries of these regions.

**Supplementary Figure 4:**
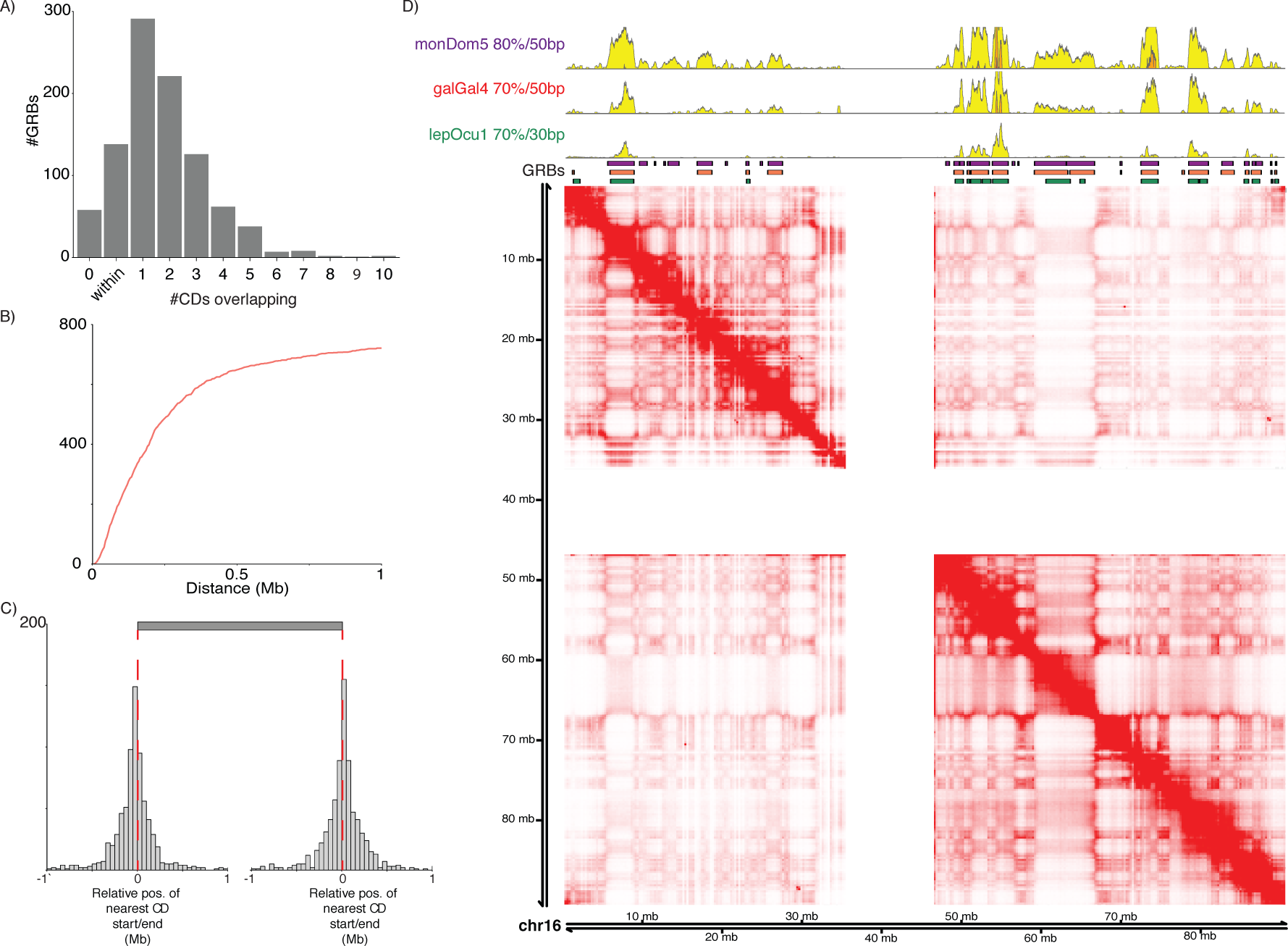
The correspondence between CNE density and topological organisation is present in high-resolution Hi-C. A) Comparison of hg19-galGal4 GRBs with a set of outermost contact domains (CD) identified in Rao *et al.* identified that a large number of GRBs were located within a CD or overlapping a single CD. The large number of GRBs which do not appear to be located within a contact domain is potentially due to the high false negative rate of the arrowhead domain finding algorithm. B) Cumulative distribution of the distance to nearest CD boundaries from GRB boundaries in GM12878 i.e. both the start and end position of a GRB lie within X kb of the nearest CD start and end. C) Relative position of the nearest CD start/end compared to the boundaries of hg19-galGal4 GRBs. D) In kilobase resolution Hi-C of GM12878 cells, the topological organisation of human chromosome 16 is highly concordant with the distribution of CNEs identified using opossum, chicken and spotted gar. Results are robust in other cell lines (data not shown).

**Supplementary Figure 5:**
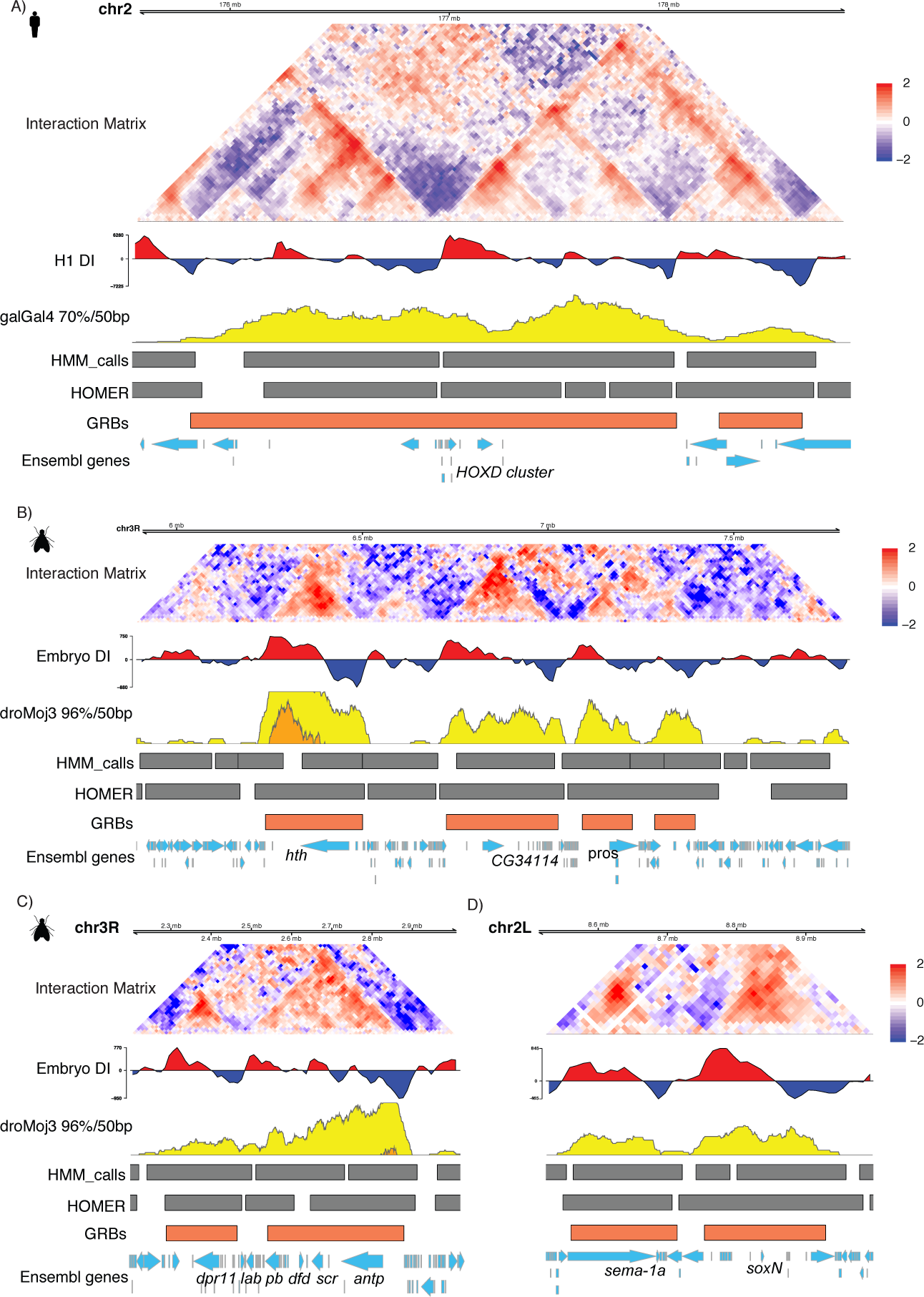
Examples of genomic regulatory blocks and their associated interaction landscapes in human and *Drosophila*. GRBs at loci in both human and *Drosophila* show strong concordance with the structure of regulatory domains proposed from Hi-C. TADs, generated using both HOMER and HMM calls, along with associated directionality index and normalised interaction matrix show a striking concordance with the boundaries of putative GRBs and with CNE density in general. A) The *HoxD* locus in human (chr2:175575640-178817020) is situated between two TADs, with its constituent genes showing interactions with regulatory elements in surrounding TADs depending on the developmental context [108, 52, 109, 110]. The proposed *HoxD* GRB recapitulates the span of known regulatory interactions better than the regulatory domains predicted by Hi-C. B) In *Drosophila*, a region spanning (chr3R:5900280-7793200) contains *hth*, *CG34114* and *pros*, all of which have important roles in development. C) The *Drosophila Antennapedia* complex (chr3R:2198960-3016900) is one of two clusters of Hox genes in fly and contains genes necessary for the proper development of the *Drosophila* body plan. D) A region (chr2L:8510000-8990000) contains two GRBs containing the transcription factor *soxN* and the secreted transmembrane protein *sema-1a*. *soxN* is the *Drosophila* homolog of the *Sox1/2/3* transcription factors and is required for the generation of neural progenitors during development, while *sema-1a* is a neuronally expressed protein involved in regulating the localisation of axons during neurogenesis.

**Supplementary Figure 6:**
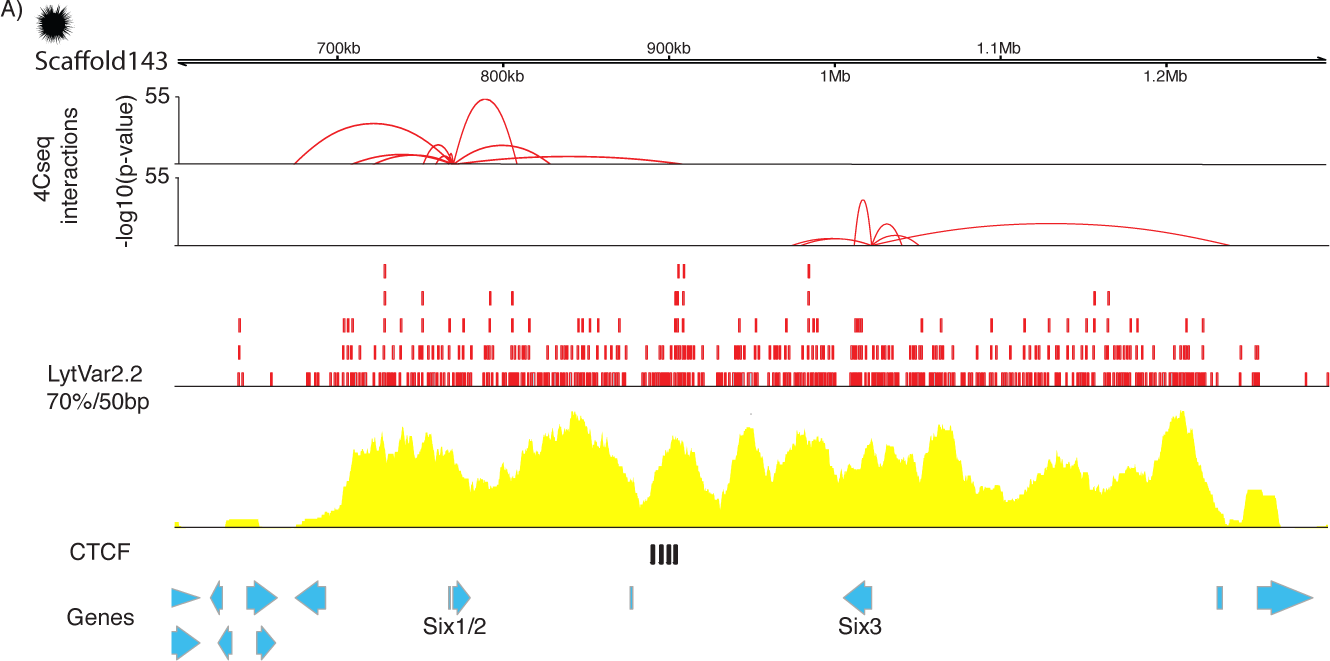
The distribution of CNEs in sea urchin correlates with the extent of interactions at the Six locus. A) The distribution of CNEs, identified in comparisons with *Lytechinus variegatus* (LytVar2.2), at the Six1/4/6 locus (Scaffold143:600000-1300000) in *S. purpuratus* correlates with the span of interactions identified in 4Cseq [111, 54]. While it is not possible to demarcate the boundary of the regulatory domains of *Six1/2* and *Six3*, there does appear to be local minima in CNE density close to the domain boundary defined by binding of CTCF. Data was obtained from Gomez-Marin *et al.* (GSE66900). Interaction tracks visualised using GenomicInteractions [112].

**Supplementary Figure 7:**
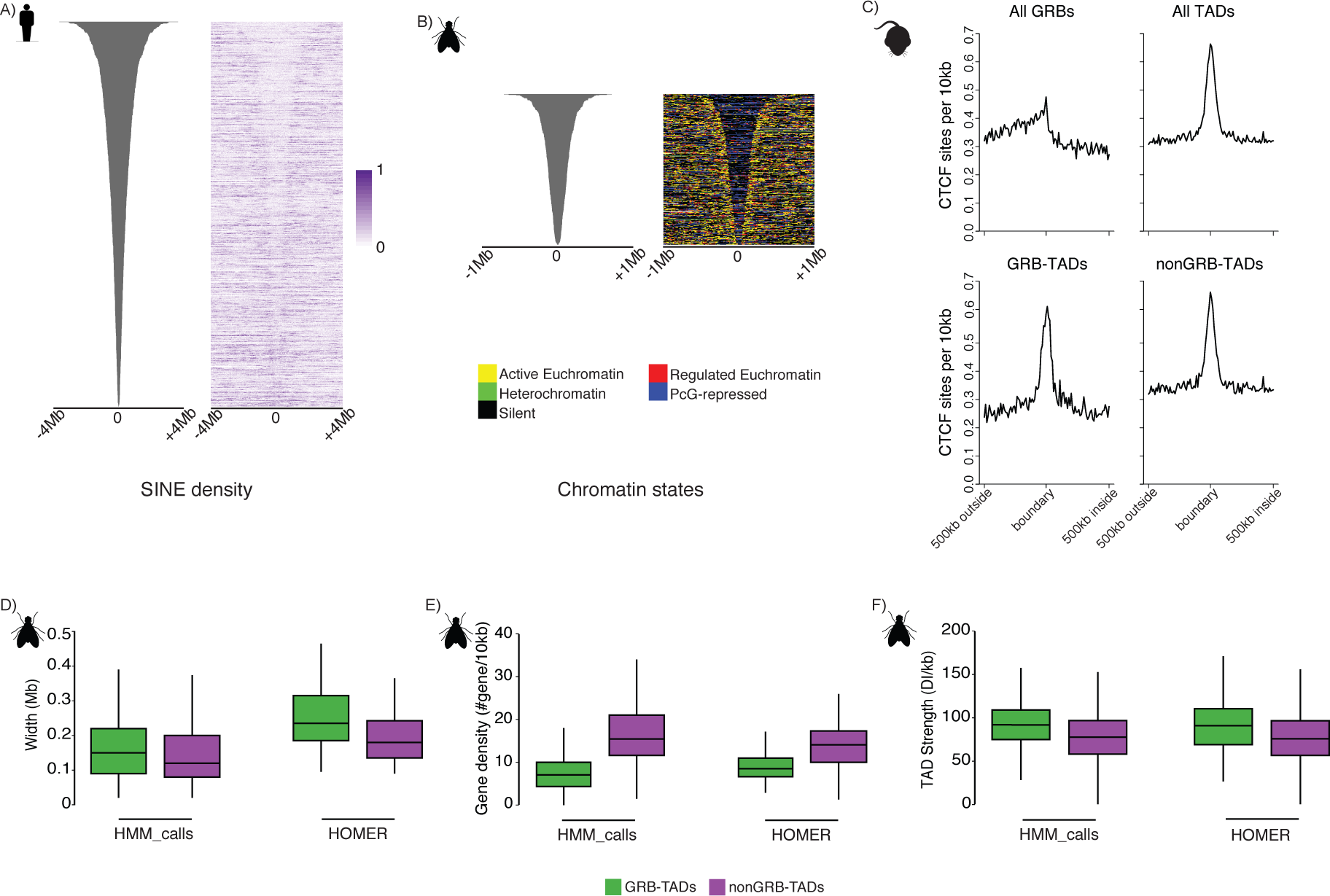
Several sets of features distinguish between TADs associated with extreme non-coding conservation (GRB-TADs) from those without (nonGRB-TADs). A) Distribution of SINE elements in a 8Mb window centred on the midpoint of hg19-galGal4 GRBs shows a clear depletion of SINEs within GRBs compared to surrounding regions. B) Chromatin states in a 2Mb window centred on the midpoint of dm3-droMoj3 GRBs. The vast majority of GRBs are covered primarily with blue and black chromatin, with a clear depletion in green and yellow chromatin. C) CTCF is enriched at GRB and TAD boundaries. CTCF sites per 10kb in a 1Mb window around GRB boundaries, TAD boundaries, and TADs separated into GRB-TADs and nonGRB-TADs (see Methods). D) The sizes of GRB-TADs identified in *Drosophila* whole embryos are significantly longer than nonGRB-TADs identified using either HOMER (median width 235kb vs. 185kb, p <0.001) or HMM calls (median width 150kb vs. 120kb, p=0.014). E) *Drosophila* embryo GRB-TADs are associated with lower protein-coding gene density than nonGRB-TADs identified using either HOMER (median # genes 8.50 vs. 14.06 p <1e-6) or HMM calls (median # genes 7.10 vs. 15.45 p <1e-6). F) Distribution of TAD strength shows that *Drosophila* GRB-TADs are significantly stronger than nonGRB-TADs identified using either HOMER (median strength 128.54 vs. 91.41, p <1e-6) or HMM calls (median strength 123.08 vs. 72.95, p <1e-6).

**Supplementary Figure 8:**
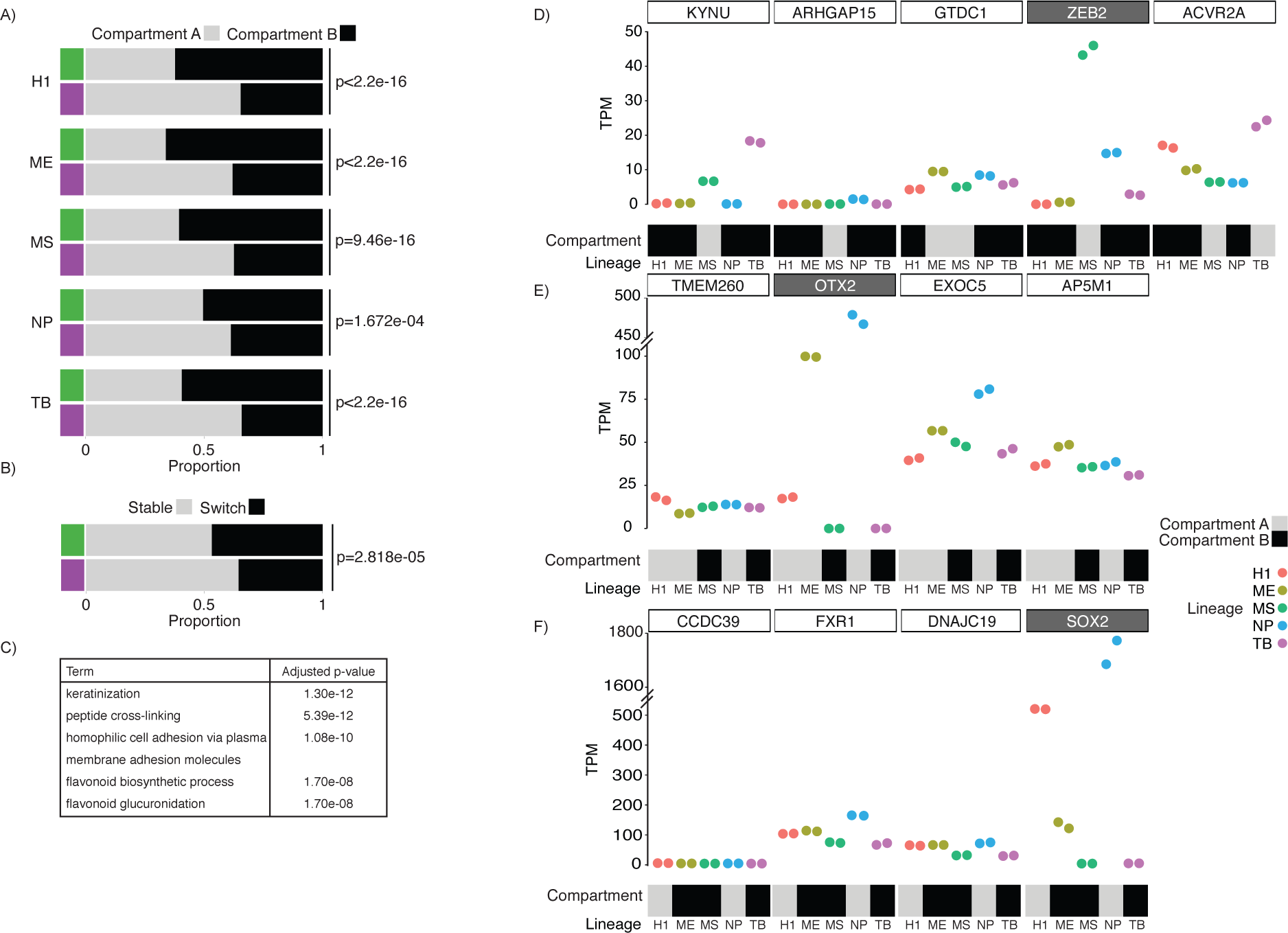
GRB-TADs are associated with compartment B, with changes in compartment between lineages associated with changes in expression of the GRB target gene. A) GRB-TADs (HMM calls) are preferentially associated with Compartment B in all of the lineages investigated. B) GRB-TADs are more likely to switch compartment at least one of the five lineages investigated (i.e. A-B or B-A) than nonGRB-TADs. C) Simplified significant GO Biological process enrichment for genes located within nonGRB-TADs, but which changed compartment in at least one of the five lineages. D) A GRB (chr2:143298358-148669176) contains the developmental regulator *ZEB2* and several bystander genes. The only gene at this locus which shows dramatic upregulation in MS, when the region is now located in Compartment A is *ZEB2*. E) The GRB (chr14:56879402-57783739) containing *OTX2*, a homeobox containing TF, is located in compartment A in ME and NP concordant with an upregulation of this gene in those lineages. Nearby bystander genes do not show any large change in their expression profiles and appear to be unaffected by compartment switching. F) A GRB (chr3:180436332-182390735) contains *SOX2*, a TF important in the maintenance of pluripotency and neurogenesis [113, 114]. The same pattern observed at the *OTX2* and *ZEB2* loci is observed with the location of this region in Compartment A in NP and H1, largely reflecting the upregulation of *SOX2* in those lineages.

**Supplementary Figure 9:**
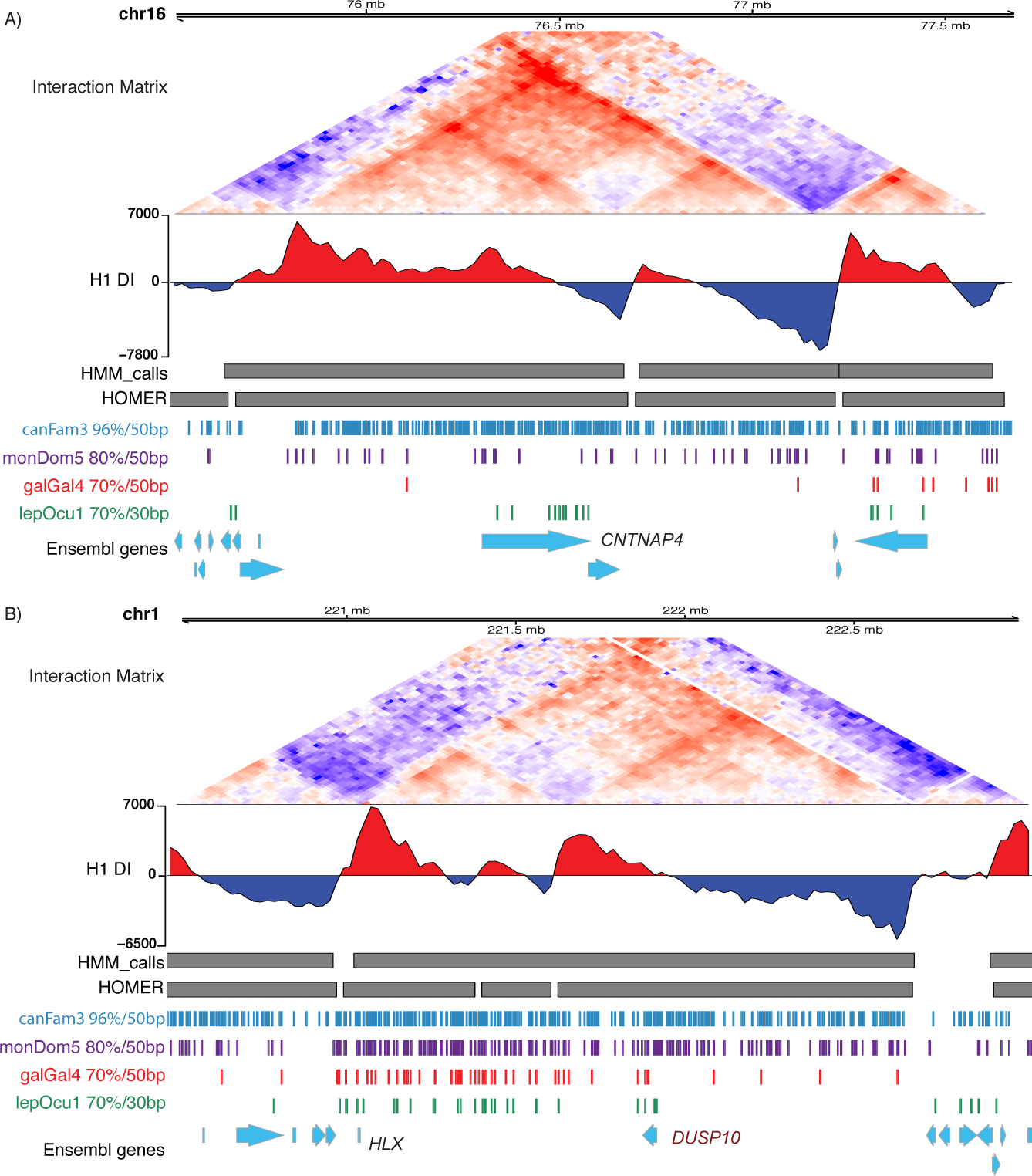
Patterns of non-coding conservation provide insights into cis-regulatory evolution at a number of loci. A) The *CNTNAP4* locus (chr16:75500000-77690000) has a limited number of CNEs identifiable in comparisons between human and spotted gar, but lacks CNEs in comparisons between human and chicken. However, comparisons between human and opossum or dog, identifies a set of CNEs, which are predictive of the topological organisation at this locus. B) A region containing the homeobox *HLX* (chr1:220500000-223000000) shows presence of CNEs in the region from *HLX* to *DUSP10*. Although there is strong conservation of the left boundary of this region, the right boundary of this locus appears to be different dependent on the species involved. CNEs may be recruited in the region to the right of *DUSP10* suggesting potential evolutionary dynamics.

**Supplementary Figure 10:**
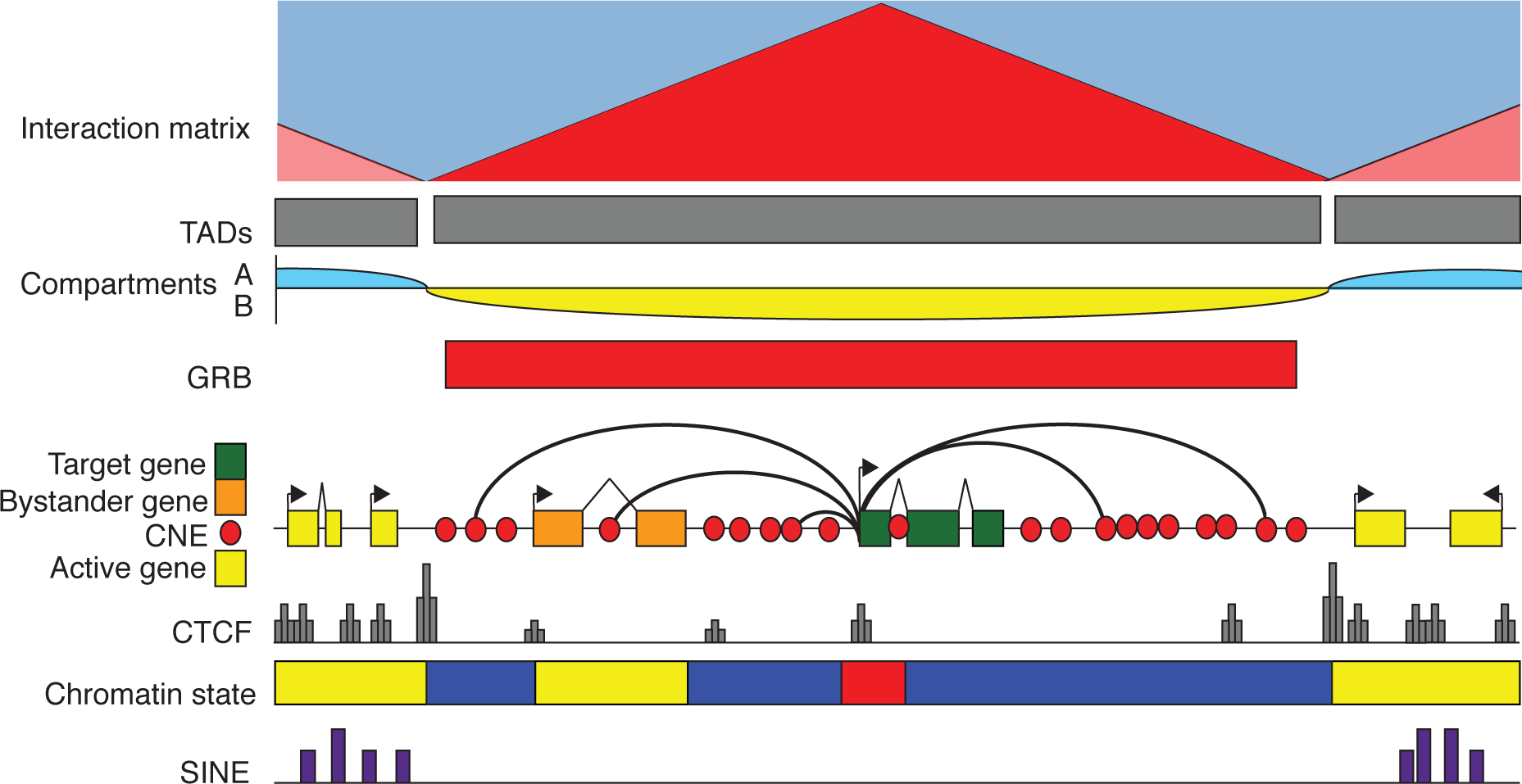
Schematic for relationship of GRB-TADs and CNEs. At loci containing important developmental regulators, the boundaries of TADs can be predicted from the distribution of CNEs. These TADs appear to be both longer and stronger than TADs lacking CNEs and are preferentially associated with compartment B. All of the regulatory elements and CNEs within this GRB-TAD are dedicated to the regulation of the GRB target gene. This regulatory domain is depleted for both CTCF and SINE elements inside it, while exhibiting enrichment for constitutive binding of CTCF at its boundaries.

